# The Fatty Acid Binding Protein Family Represents a Novel Target in Multiple Myeloma

**DOI:** 10.1101/2022.07.01.498411

**Authors:** Mariah Farrell, Heather Fairfield, Michelle Karam, Anastasia D’Amico, Connor S. Murphy, Carolyne Falank, Romanos Sklavenitis Pistofidis, Amanda Cao, Catherine R. Marinac, Julie A. Dragon, Lauren McGuinness, Carlos Gartner, Reagan Di Iorio, Edward Jachimowicz, Victoria DeMambro, Calvin Vary, Michaela R. Reagan

**Affiliations:** Center for Molecular Medicine, Maine Health Institute for Research, Scarborough, ME, USA; Graduate School of Biomedical Science and Engineering, University of Maine, Orono, ME, USA; Tufts University School of Medicine, Boston MA, USA; Dana-Farber Cancer Institute, Boston, MA, USA; Harvard Medical School, Boston, MA, USA; University of Vermont, Burlington, VT, USA; University of New England, Biddeford, ME, USA

**Keywords:** Myeloma, fatty acid binding protein, FABP, FABP5, RNA-Seq, Proteomics

## Abstract

**Background:** Multiple myeloma is an incurable plasma cell malignancy with only a 53% 5-year survival rate, highlighting a critical need for new multiple myeloma vulnerabilities and therapeutic avenues. Herein, we explored a novel multiple myeloma target: the fatty acid binding protein (FABP) family.

**Methods:** Myeloma cells treated with FABP inhibitors (BMS3094013 and SBFI-26) were examined *in vivo* and *in vitro* for cell cycle, proliferation, apoptosis, mitochondrial membrane potential, cellular metabolism (oxygen consumption rates and fatty acid oxidation), and DNA methylation. Myeloma cell responses to BMS309403 and/or SBFI-26 were assessed with RNA-sequencing and proteomic analysis, and confirmed with western blotting and qRT-PCR. Myeloma cell dependency on FABPs was assessed using DepMap. Finally, MM patient datasets (CoMMpass and GEO) were mined for *FABP* expression correlations with clinical outcomes.

**Results:** Myeloma cells treated with FABPi or with *FABP5* knockout (generated via CRISPR/Cas9 editing) exhibited diminished proliferation *in vitro*. FABPi had potent anti-tumor effects both *in vitro* and *in vivo* in two pre-clinical MM mouse models where increased mouse survival was observed. FABPi negatively impacted mitochondrial respiration and reduced expression of MYC and other key signaling pathways in MM cells. Clinical data demonstrated worse overall and progression-free survival in patients with high *FABP5*.

**Conclusions:** This study establishes the FABP family as a therapeutically actionable dependency in multiple myeloma with a multitude of actions and cellular roles that result in the support of myeloma progression.

**Statement of translational relevance:** Multiple myeloma (MM) is an incurable disease of the plasma cell and MM patients require better treatments as soon as possible. The fatty acid binding protein (FABP) family plays a number of roles in cells, including supporting fatty acid oxidation, lipid shuttling and signal transduction. Here, we demonstrate with CoMMpass and other clinical data that FABPs represent a biomarker for aggressive disease in MM, and are a novel, targetable protein family expressed by myeloma cells. Pharmacologically inhibiting FABPs kills tumor cells and induces cell cycle arrest *in vitro* and in pre-clinical models. Mechanisms of action are multitudinous, as we discovered with RNA-sequencing, proteomic analysis, and phenotyping assays. Cell metabolism, cell signaling, cell stress, and epigenetic signatures were altered in MM cells when FABPs were inhibited. In summary, targeting FABP5 holds great therapeutic potential for killing diseased cells, with few negative off-target effects on healthy cells.

## INTRODUCTION

Fatty acid binding protein (FABP) family members are small (12-15 kDa) proteins that reversibly bind lipids [1, 2]. The ten human FABP isoforms are functionally and spatially diverse, consisting of ten anti-parallel beta sheets, which form a beta barrel that shuttles fatty acids across membranes of organelles including peroxisomes, mitochondria, nuclei, and the endoplasmic reticulum [3]. FABPs influence cell structure, intracellular and extracellular signaling, metabolic and inflammatory pathways [2], and maintain mitochondrial function [4]. While most cell types express a single FABP isoform, some co-express multiple FABPs that can functionally compensate for each other if needed [5, 6], suggesting that broad FABP targeting may be necessary. FABP insufficiencies in humans and mice induce health benefits (eg. protection from cardiovascular disease, atherosclerosis, and obesity-induced type 2 diabetes), suggesting these to be safe therapeutic targets [7–9].

Multiple myeloma (MM), a clonal expansion of malignant plasma cells, accounts for ∼10% of hematological neoplasms [10]. Myeloma cell growth initiates in and spreads throughout the bone marrow, leading to aberrant growth and destruction of the bone and bone marrow [11]. Treatments for myeloma patients have greatly improved within the past two decades [1], but most patients eventually relapse, demonstrating the need to pursue more novel MM treatments. Few therapies are designed to specifically target molecules involved in the unique metabolism of MM cells despite recent findings that MM cells uptake fatty acids through fatty acid transport proteins, which can enhance their proliferation [12]; as such, we sought to investigate a potential role of the FABPs in myeloma.

Links between FABP4 and cancer have been demonstrated in prostate, breast, and ovarian cancer, and acute myeloid leukemia (AML) [13–20]. FABP5 has been less widely studied in cancer, but is known to transport ligands to PPARD [21] which can intersect with many pro-tumor pathways by increasing proliferative, survival [22–24]), and angiogenic factors [25]), and decreasing tumor suppressor expression [23]. Herein we explored the oncogenic function of the FABPs in MM by examining therapeutic targeting with FABP inhibitors (FABPi) in multiple cell lines, genetic knockout of *FABP5,* pre-clinical models, large cell line datasets, and multiple patient datasets. Our results suggest FABPs are a novel target in MM due to the plethora of important biological functions that FABPs modulate to control cellular processes at multiple levels.

## RESULTS

### *FABP5* is vital for MM cells and genetic knockout results in reduced cell number

We first examined FABP gene expression in MM cell lines and found that *FABP5* was the most highly-expressed FABP in GFP^+^/Luc^+^MM.1S and RPMI-8226 cells (Additional File 1, Supplemental Table 1, [26]) and that other FABPs were also expressed to a lesser extent (eg. FABP3, FABP4 and FABP6). FABP5 protein was also robustly expressed in these cells (Supplemental Fig. 1). We then defined the landscape of FABP vulnerabilities in myeloma cells using the Broad Institute’s Cancer Dependency Map (DepMap) [27]. Of all the FABPs, only *FABP5* exhibited a negative CERES Score (-0.30) in all 20 MM cell lines, demonstrating a strong reliance on *FABP5* for their survival (Supplemental Fig. 2A). Interestingly, all cancer types within the DepMap database had negative *FABP5* CERES values (Supplemental Fig. 2B). However, since FABPs were not pan-essential (CERES score of ≤-1), targeting these should not present toxicity risks. Importantly, many fatty acid metabolism genes, including *FABP5*, had negative CERES scores (shown in blue) in MM cells (Supplemental Fig 2C).

**Figure 1.**
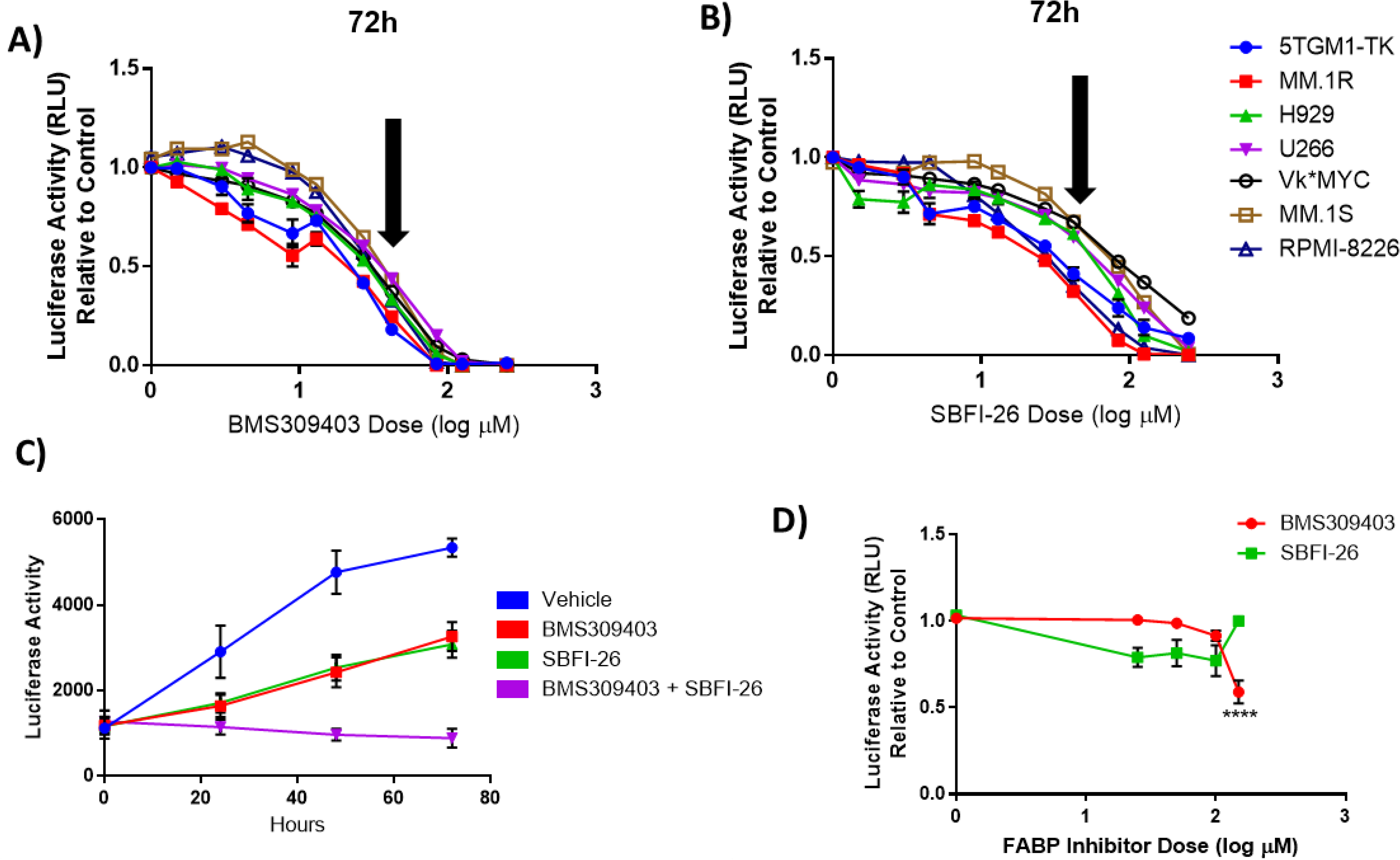
FABPi significantly impair MM cell growth and induces apoptosis. MM cell lines cell number after being exposed to (A) BMS309403 and (B) SBFI-26 for 72 hours; 50 µM dose (∼EC50) indicated by arrows. C) GFP^+^/Luc^+^MM.1S cell numbers when treated with inhibitors in combination (50 µM each). MM.1S vs BMS309403 (24 hrs, *; 48 hrs, ****; 72 hrs, ****). MM.1S vs SBFI-26 (24 hrs, *; 48 hrs, ****; 72 hrs, ****). MM.1S vs BMS309403 + SBFI-26 (24 hrs, ***; 48 hrs, ****; 72 hrs, ****). BMS309403 vs BMS309403 + SBFI-26 (48 hrs, **; 72 hrs, ****). SBFI-26 vs BMS309403 + SBFI-26 (48 hrs, **; 72 hrs, ****). Two-way ANOVA analysis with Tukey’s multiple comparisons test analysis. D) CellTiter-Glo analysis of Human mesenchymal stem cell number after being exposed to BMS309403 and SBFI-26 for 72 hours. Data are mean ± SEM unless otherwise stated and represent averages or representative run of at least 3 experimental repeats. One-way ANOVA with Dunnett’s multiple comparison test significance shown as *p < 0.05. **p < 0.01. ***p<0.001. ****p < 0.0001. **** *P* < 0.0001.

**Figure 2.**
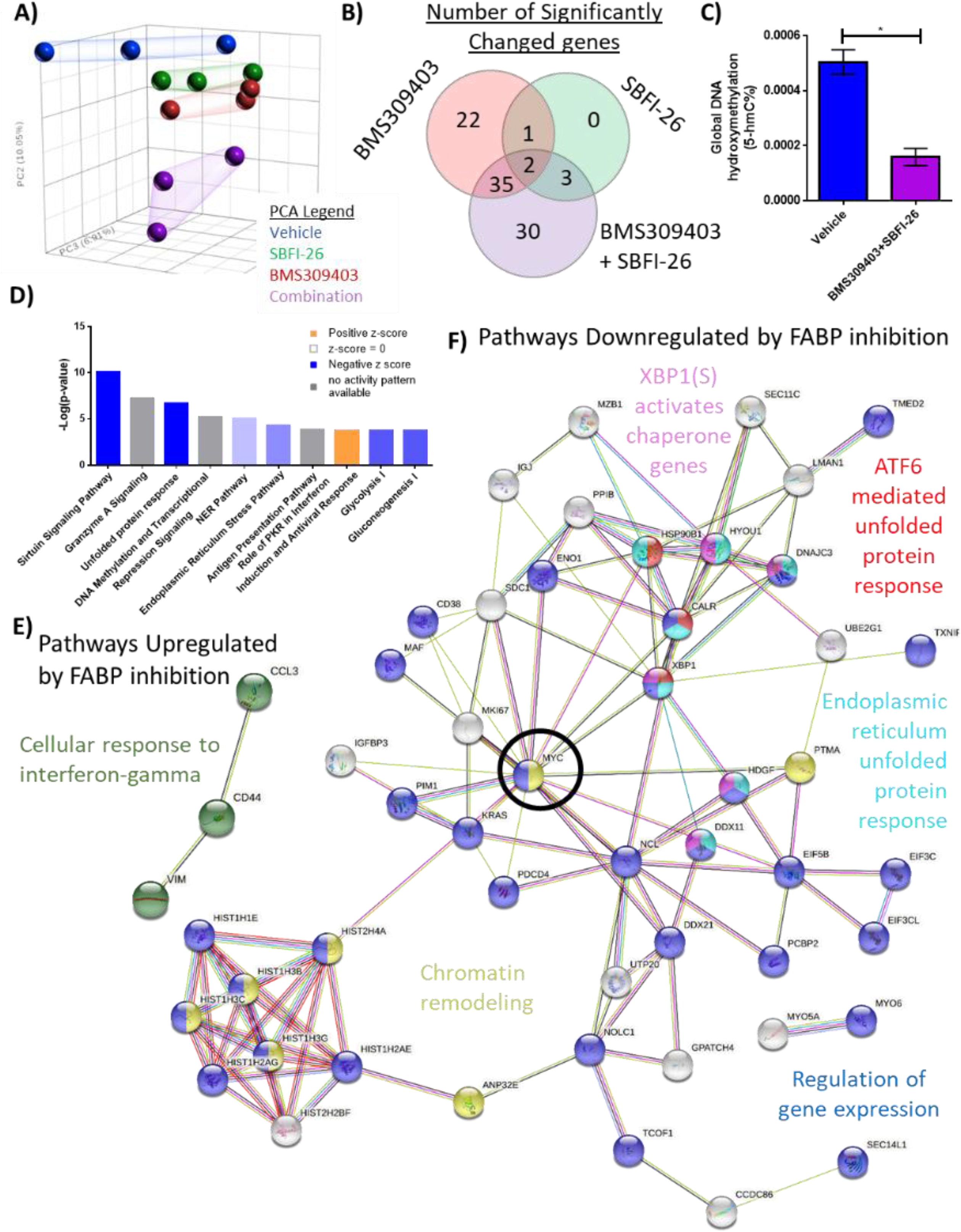
RNA sequencing analysis of MM1S cells treated with FABPi for reveals unique gene expression patterns. A) Principal component analysis of cells after 24 hour treatments. Venn diagram displays the overlapping and specific genes dysregulated with FABPi (FDR cutoff of 0.2). C) Global hydroxymethylation DNA analysis of MM.1S cells after 24 hours of combination treatment. D) Ingenuity pathway analysis of RNA-Seq results (p-value of overlap by Fisher’s exact test, significance threshold value of p<0.05(-log value of 1.3)). Stringdb (FDR cutoff of 0.2) of the combination therapy versus control showing (E) the 1 upregulated pathway and (F) 5 of the many downregulated pathways. MYC, a central node, is circled for emphasis.

As such, we next examined the effect of *FABP5* knockout (KO) in MM cells, as this was the most highly expressed FABP in all three cell lines. *FABP5* mutant (*FABP5*^MUT^) MM.1R cells exhibited a 94% editing efficiency with a ∼59% KO efficiency after expansion (Supplemental Fig. 3A, B). We observed an 84% reduction in *FABP5* expression in the edited pool (Supplemental Fig. 3C), confirming functional *FABP5* knockdown. *FABP4* expression was not altered (Supplemental Fig. 3D), but *FABP6* expression was increased in the edited cells (Supplemental Fig. 3E)*. FABP5* edited cells had slight, but significantly reduced cell numbers at 48, 72, and 96 hours versus controls (Supplemental Fig. 3F).

**Figure 3.**
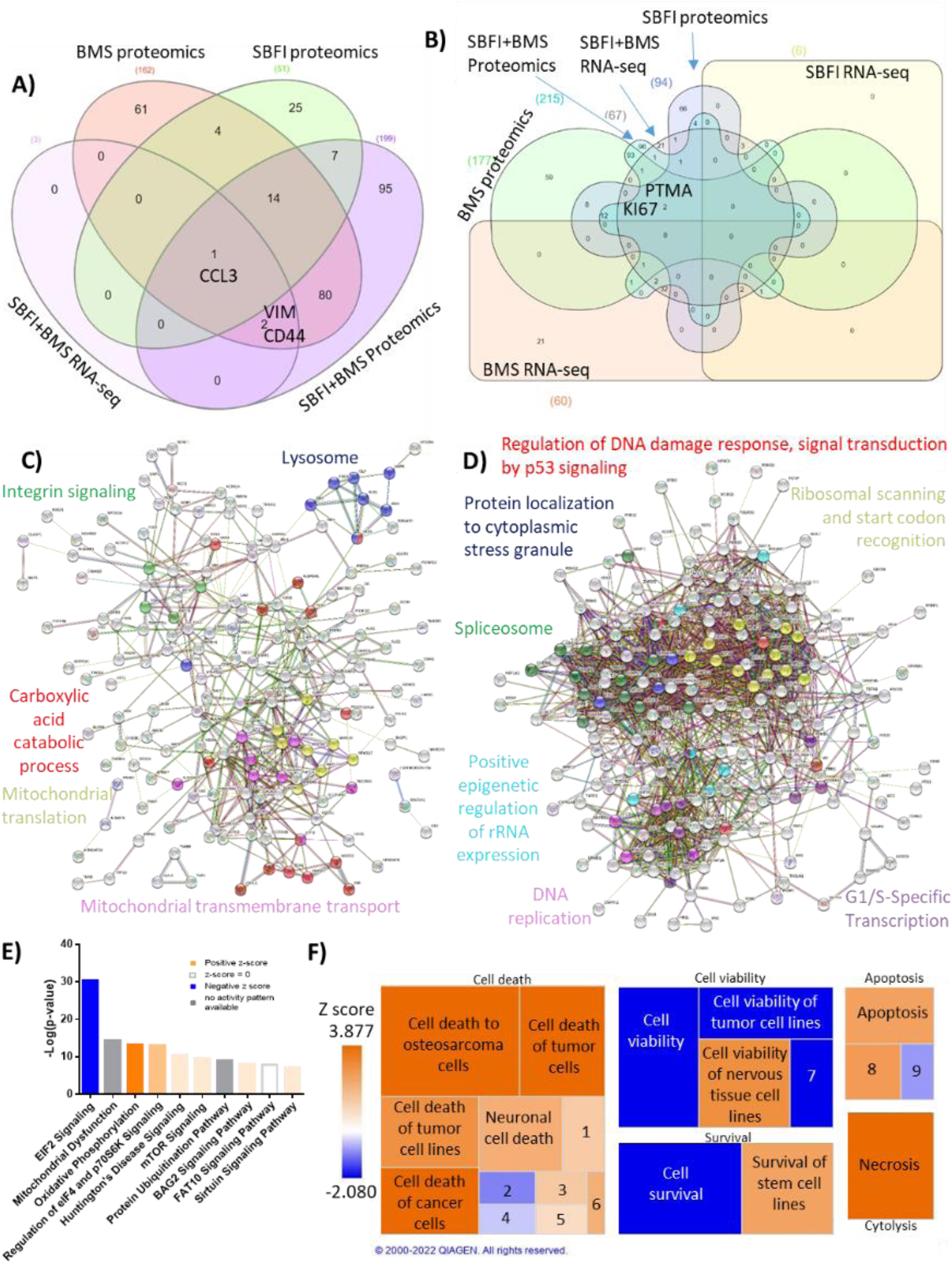
48 hour proteomic analysis of MM1S cells treated with FABPi reveals a unique protein signature. MM.1S cells were assessed by proteomics after 48 hour treatments with BMS309403 (50 µM), SBFI-26 (50 µM) or the combination, and compared to results from RNA-Seq. N=3 biological replicates and 3 technical replicates Venn diagram comparison of (A) upregulated genes and (B) downregulated proteins in proteomics and RNA sequencing among BMS309403 and SBFI-26 treated cells compared to vehicle. C-F) Pathway analysis of proteomic data of significantly upregulated or downregulated proteins in MM.1S cells treated with both FABPi (BMS309403+SBFI-26). C, D) String analysis of upregulated (C) or downregulated (D) pathways. E) Top 10 significantly changed pathways with FABP inhibition. For IPA analysis, orange represents positive z-score, blue indicates a negative z-score, gray represents no activity pattern detected and white represents a z-score of 0. F) Ingenuity pathway analysis of the Cell Death and Survival heatmap. Numbers in boxes represent: 1) Cell death of melanoma lines; 2) Cell death of carcinoma cell lines; 3) Cell death of neuroblastoma cell lines; 4) Cell death of breast cancer cell lines; 5) Cell death of connective tissue cells; 6) Cell death of fibroblast cell lines; 7) Cell viability of myeloma cell lines; 8) Apoptosis of tumor cell lines; 9) Apoptosis of carcinoma cell lines.

### Pharmacological inhibition of FABPs reduces myeloma cell proliferation *in vitro*

Having observed potential compensation among FABP family members in the *FABP5*^MUT^ cells, we used two well-known FABP inhibitors (FABPi): BMS309403 and SBFI-26, which specifically and potently inhibit FABPs by binding their canonical ligand-binding pockets. Ligand-binding assays determined that BMS309403 has Ki values in solution of <2, 250 and 350 nM for FABP4, FABP3 and FABP5, respectively, and SBFI-26 has Ki values of 900 and 400 nM for FABP5 and FABP7 respectively, as reported on the manufacturers’ datasheets [28]. A 72-hour dose curve of either BMS309403 or SBFI-26 demonstrated a dose-dependent decrease in cell number in all myeloma lines tested (Fig. 1A, B; Additional File 1, Supplemental Table 2, 3). Similar effects were seen at earlier time points and in other myeloma cell lines (Supplemental Fig. 4). 50 µM of BMS309403, SBFI-26, or the combination (50 µM BMS309403 + 50 µM SBFI-26) significantly reduced cell numbers at 24, 48, and 72 hours by 39%, 42%, and 83% respectively in GFP^+^/Luc^+^MM.1S cells (Fig. 1C), suggesting the ability to synergize FABPi.

**Figure 4.**
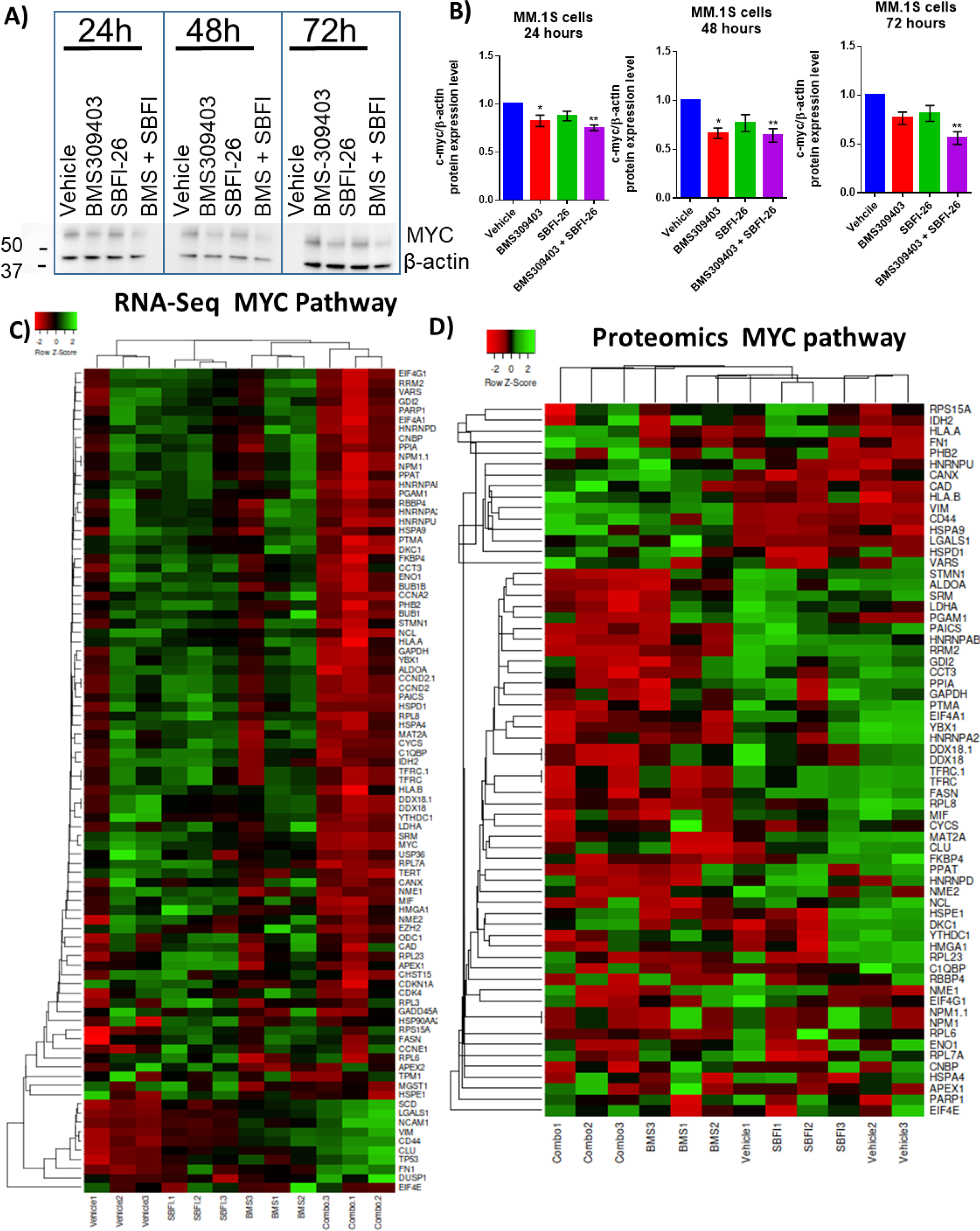
FABPi targets MYC and the MYC pathway. A) Representative western blot and B) quantification of MYC protein and β-actin (housekeeping control) at 24, 48, and 72 hours after treatment with BMS309403 (50 µM), SBFI-26 (50 µM), or the combination. Data represent mean ± SEM from n=3 biological repeats, analyzed with one-way ANOVA with significance shown as *p<0.05. **p<0.01. ****p<0.0001. C) RNA-seq and D) Proteomic analysis of expression of genes/proteins involved in MYC signaling shown as heatmap visualizations. Curated lists are based on IPA MYC Pathway list, known MYC-regulated genes, and proteins present in proteomics.

Non-cancerous cells were found to be much less sensitive to FABPi (Fig. 1D), supporting prior literature showing the safety of these inhibitors [13, 17]. No change in amount or localization of FABP5 protein after treatment with FABPi in GFP^+^/Luc^+^MM.1S or RPMI-8226 cells was observed either by immunofluorescence (Supplemental Fig. 5) at 24 hours, or by Western blotting at 24, 48, or 72 hours in GFP^+^/Luc^+^MM.1S cells (Supplemental Fig. 6A, B). These data suggest that FABP activity, but not protein expression, was decreased by the inhibitors. Conversly, recombinant FABP4 and FABP5 did not affect MM.1S cell number (Supplemental Fig. 7A, B).

**Figure 5.**
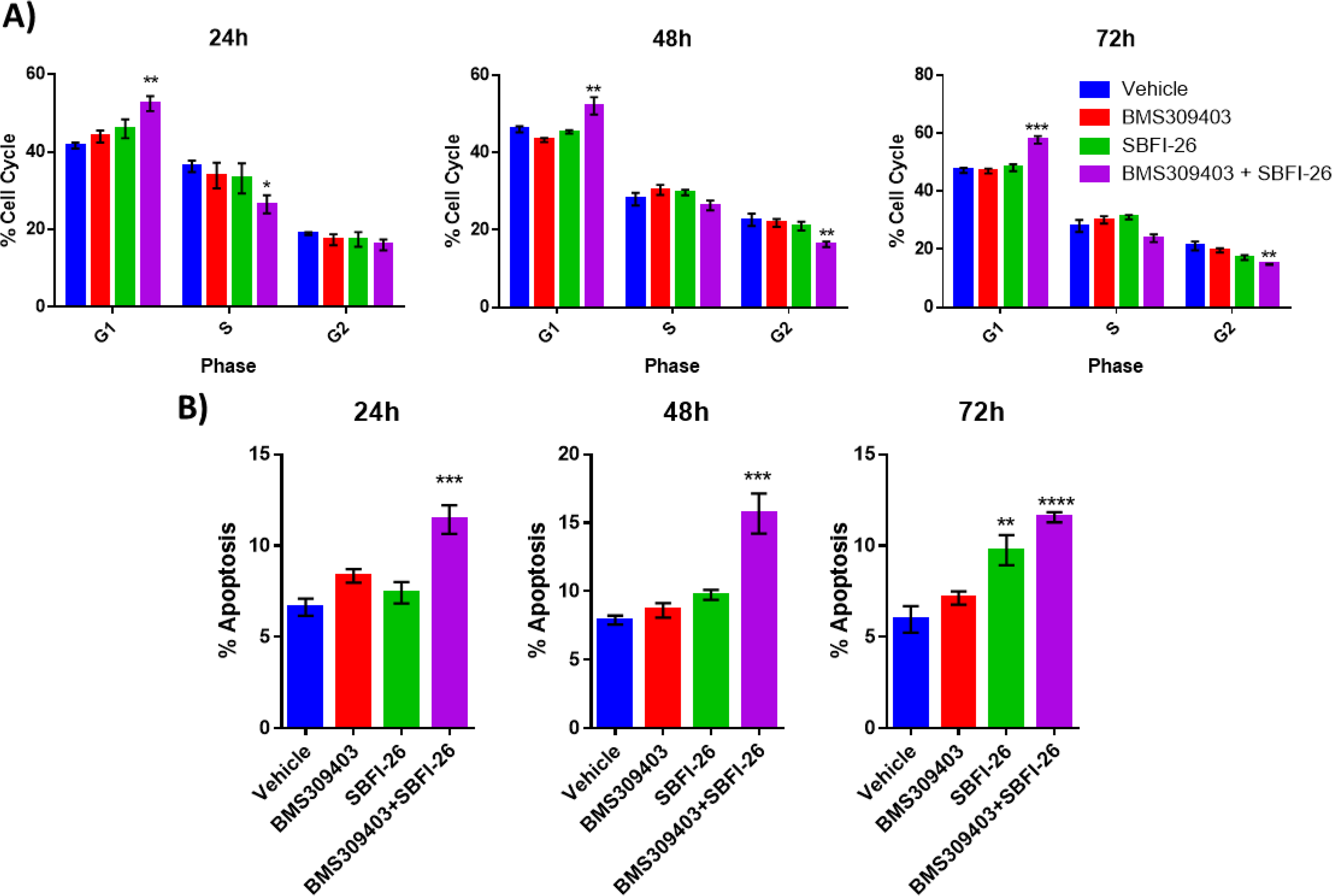
FABPi significantly impair MM cell growth and induces apoptosis. A) GFP+/Luc+MM.1S cell cycle states with the FABPi in combination (50 µM each). B) Apoptosis in MM.1S cells with FABPi. Data are mean ± SEM unless otherwise stated and represent averages or representative run of at least 3 experimental repeats. One-way ANOVA with Dunnett’s multiple comparison test significance shown as *p < 0.05. **p < 0.01. ***p<0.001. ****p < 0.0001.

**Figure 6.**
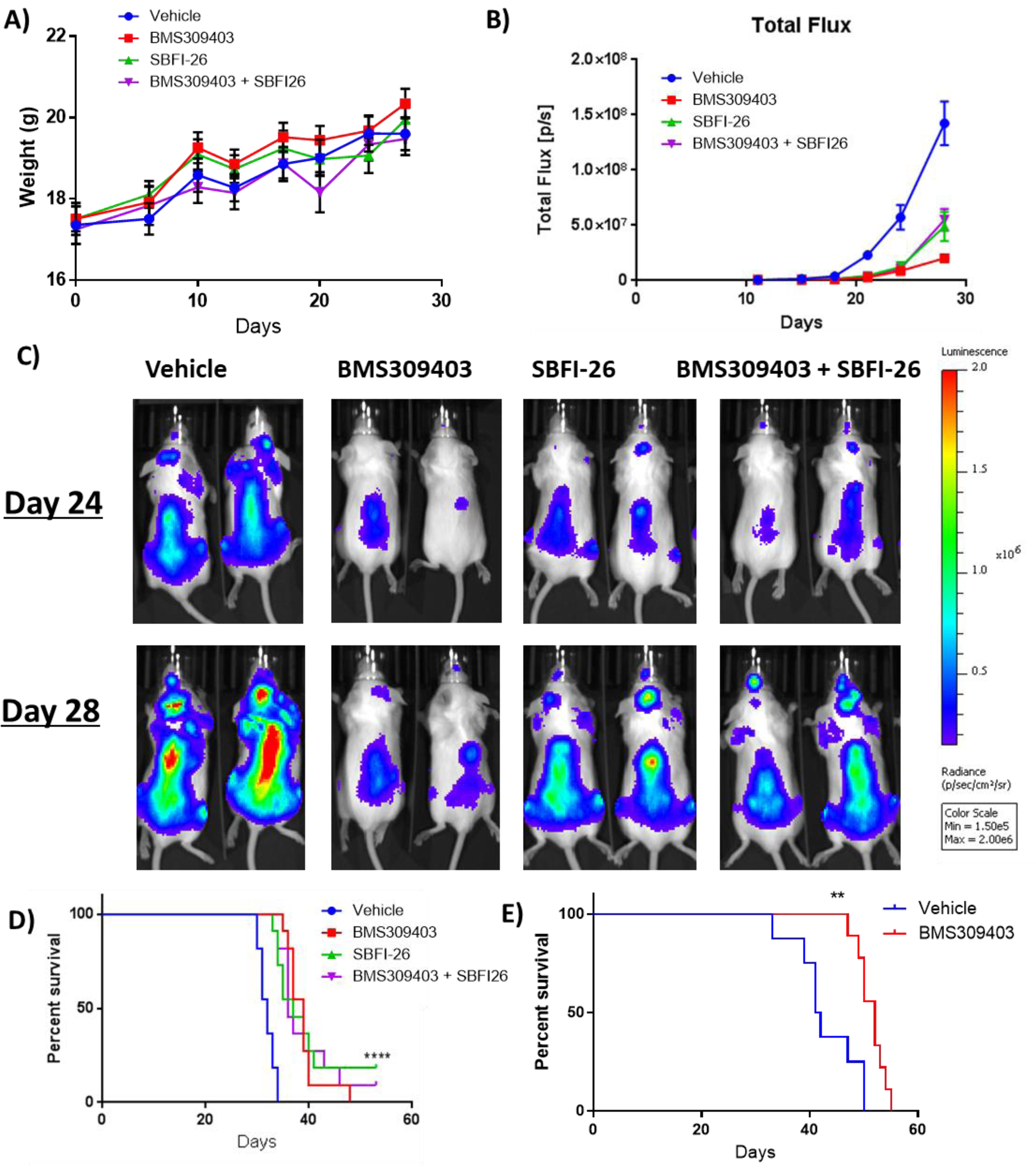
FABPi extend survival and decrease tumor burden in myeloma xenograft and syngeneic mouse model. A) Mouse weights normalized to day 0 for each group (treated with BMS309403, SBFI-26, or the combination) from day of injection plotted as Mean ± SEM. B) Tumor burden assessed by bioluminescence imaging (BLI) in MM.1S model. One-way ANOVA with Dunnett’s multiple comparison test significance shown as *p < 0.05. **p < 0.01. ***p<0.001. ****p < 0.0001. Vehicle vs BMS309403 (24 days, ****; 28 days, ****). Vehicle vs SBFI-26 (24 days ****; 28 days, ****). Vehicle vs BMS309403 + SBFI-26 (24 hrs, ****; 28 days, ****). BMS309403 vs BMS309403 + SBFI-26 (24 days NS; 28 days, ***). SBFI-26 vs BMS309403 + SBFI-26 (24 and 28 days, NS). BMS309403 vs SBFI-26 (24 hrs, NS 28 days, **). C) Representative BLI images at days 24 and 28 from B. D) Survival of mice mice from B; analysis performed by Kaplan-Meier Survival Analysis, Log-Rank (Mantel-Cox) test, p <0.0001, n=11. E) Survival of KaLwRij mice injected with 5TGM1 cells. Survival analysis performed by Kaplan-Meier Survival Analysis, Log-Rank (Mantel-Cox) test, p=0.0023, Vehicle n=8, BMS309403 n=9.

### FABPi induce gene expression changes in MM cells that affect a range of cellular processes and pathways linked to survival

To identify transcriptional changes that may mediate the effects of FABP inhibition on cell number, we treated GFP^+^/Luc^+^MM.1S cells with the single FABPi (50 µM) or the combination (50 µM of each) for 24 hour, isolated total RNA, and performed RNA sequencing. Principal component analysis (PCA) demonstrated that the FABPi groups exhibited distinct gene expression profiles, and that the combination treatment differed the most from vehicle-treated cells (Fig. 2A). Over 14,000 genes were analyzed, revealing 93 significant differentially expressed (DE) genes within all three treatment groups, compared to the vehicle control (FDR<0.2): 90 downregulated and 3 upregulated (Fig. 2B; Additional File 1, Supplemental Table 4). Consistent with decreased levels of transcription, we also observed significantly lower levels of 5-hydroxymethylcytosine in FABPi-treated cells compared to vehicle-treated cells (Fig. 2C), suggesting decreases in active chromatin. This finding is consistent with previous reports linking FABP depletion to DNA methylation signatures in other cancers [13, 18].

To further understand the mechanisms of action of FABPi, we investigated which pathways were impacted in our RNA-Seq data using STRINGdb and IPA (ingenuity pathway analysis). IPA was specifically used to investigate canonical pathways, while STRINGdb was used to examine connectivity of DE genes and enrichment for specific gene ontology terms, as well as molecules in Reactome and KEGG pathways. In total, 15 IPA canonical pathways were commonly dysregulated in all three treatment groups including Cell Cycle: G2/M DNA Damage Checkpoint Regulation, EIF2 Signaling, Sirtuin Signaling Pathway, and the NER pathway (Additional File 1, Supplemental Table 5; Fig. 2D). The one upregulated pathway according to STRING was “cellular response to interferon gamma signaling” in the combination group (Fig. 2E; Additional File 1, Supplemental Table 6). The top downregulated pathways in the combination treatment by STRING analysis are in Additional File 1, Supplemental Table 7.

Interestingly, both IPA and STRING databases revealed commonly downregulated pathways related to the unfolded protein response (UPR) and ER stress responses for BMS309403 (Supplemental Fig. 8A-C; ER Stress –log(p-value=3.78E+00)), SBFI-26 (Supplemental Fig. 8A-C), and the combination (Fig. 2D, F; Supplemental Fig. 10A). Three of the five downregulated Reactome pathways in the combination group were related to UPR or ER stress (Fig. 2F), driven by molecular players such as *XBP1*, *BIP*, and *IRE1*. Downregulation of total *XBP1* by the combination treatment was confirmed after 24 hours (Supplemental Fig. 10B) and heatmaps visually demonstrated the downregulation of genes involved in XBP1 signaling (Supplemental Fig. 10C) and the UPR (Supplemental Fig. 10D) as determined by IPA. Interestingly, *MYC*, a known oncogene, was found as a central node in STRING analysis (Fig. 2F) and in the top 10 most downregulated genes in RNA-Seq from combination treatments (Additional File 1, Supplemental Table 8).

### FABPi induces protein changes in MM cells that affect a range of cellular processes and pathways linked to survival

To identify protein changes resulting from FABPi, we treated GFP^+^/Luc^+^MM.1S cells with the single inhibitors (50 µM) or the combination (50 µM of each) for 48 hour, isolated total cell lysate proteins, and performed a mass spectrometry-based proteomic analysis. (Numbers of significant proteins, Additional File 1, Supplemental Table 9; gene names, Additional File 1, Supplemental Tables 10-15). PCA analysis showed a tight grouping of samples (Supplemental Fig. 11A); 15 genes were commonly upregulated and 15 commonly downregulated genes between all treatments (Supplemental Fig. 11B, C; Additional File 1, Supplemental Table 16, 17).

We then compared significant genes and proteins identified by both RNA-Seq and proteomics (Fig. 3A, B). CCL3, a chemokine for monocytes, macrophages, and neutrophils, was upregulated by SBFI-26, BMS309403, and their combination in proteomics, and upregulated by the combination treatments in RNA-Seq. Ki67, a proliferation marker, and PTMA, a negative regulator or apoptosis, were both significantly downregulated in the combination treatment in RNA-Seq and proteomics, and in the single drug treatments in proteomics (Fig. 3B), indicating cell death and cell cycle arrest likely result from FABPi.

STRING analysis of proteomics data suggested many other systemic changes (eg, downregulation of DNA replication and other viability/proliferation processes and upregulation of lysosome, carboxylic acid catabolic process, and mitochondrial pathways) induced by the FABPi combination treatments (Fig. 3C, D) and interesting up- and downregulated pathways by BMS309403 or SBFI-26 treatments alone (Supplemental Figs. 12A-B, 13 A-B ). IPA analysis revealed “EIF2 Signaling” to have the highest negative Z-score for all FABPi treatments in proteomics (Fig. 3E; Supplemental Figs. 12C, 13C). IPA “Cell Death and Survival” heatmap analysis showed increases in cell death and apoptosis pathways and decreases in cell viability pathways after FABPi combination treatment (Fig. 3F; Supplemental Figs. 12D, 13D). Interestingly, MYC was the most significant predicted upstream regulator, found to be strongly inhibited in the BMS309403, SBFI-26, and combination treatments from IPA proteomic analysis (Additional File 1, Supplemental Tables 18-20).

Since *MYC* was found as a central node or commonly downregulated gene/pathway in our RNA-Seq and proteomic data analyses, we investigated MYC’s role in FABP signaling in myeloma cells. We confirmed decreased *MYC* expression in GFP^+^/Luc^+^MM.1S cells treated with the FABPi combination, and also saw a trend for this in 5TGM1-TK cells treated with either inhibitor alone or the combination (Supplemental Fig. 14A, B). MYC protein level was also decreased in GFP^+^/Luc^+^MM.1S cells at 24, 48, and 72 hours with FABPi (Fig. 4A, B). MYC-regulated genes were also decreased with FABPi in both the RNA-Seq (Fig. 4C; Supplemental Fig. 9D) and proteomic data (Fig. 4D) by heatmap analysis. In RNA-Seq data, treatment with BMS309403 induced aberrant gene expression of 171 genes known to be regulated by MYC (Additional File 1, Supplemental Table 21), with 138 of those having expression patterns consistent with MYC inhibition. Similarly, co-treatment induced changes in 91 genes modulated by MYC (Supplemental Fig. 14C; 68 consistent with MYC downregulation), while 29 MYC targets were aberrantly expressed with SBFI-26 treatment (Supplemental Fig. 9D; 18 consistent with MYC downregulation).

### FABPi impair MM cell metabolism, mitochondrial function and cell viability

Having observed effects of the inhibitors on metabolic processes such as mitochondrial function and and oxidative phosphorylation in our proteomic data, we next assessed mitochondrial function and metabolic changes using a Cell Mito Stress Test (Supplemental Fig. 15A). After 24 hour treatments, all FABPi treatments decreased basal mitochondrial oxygen consumption rates (OCR) and OCR dedicated to ATP production (Supplemental Fig. 15B). Maximal respiration and spare respiratory capacity was decreased with SBFI-26 and combination treatments, suggesting FABP inhibition reduces the ability of MM cells to meet their energetic demands.

To determine the effects of FABPi on fatty acid oxidation (FAO) specifically, we treated tumor cells with etoxomir, an FAO inhibitor, with or without the FABPi (Supplemental Fig 15 C-E). The combination of FABPi alone again strongly reduced mitochondrial respiration in most of the parameters assessed in a mitochondrial stress test.

Interestingly, etoxomir treatment caused a slight, but significant reduction in OCR when it was administered, demonstrating some reliance of MM cells on FAO for mitochondrial respiration. However, the FABPi had a much greater effect on MM mitochondrial respiration than etoxomir alone, suggesting that FABPi treatment is inhibiting mitochondrial respiration through another mechanism. Also, since maximal respiration was decreased in the Etox+FABPi combination compared to FABPi alone, it is also clear that FABPi treatment does not completely block FAO when used alone. Overall, the data demonstrate that mitochondrial respiration is inhibited by FABPi. To assess whether metabolic dysfunction could be caused by damaged mitochondria, we utilized tetramethylrhodamine, ethyl ester (TMRE) staining and flow cytometric analysis. GFP^+^/Luc^+^MM.1S cells treated with BMS309403 or the combination (BMS309403 +SBFI-26) had decreased TMRE staining (Supplemental Fig. 16A, B), suggesting that BMS309403 damages MM cell mitochondria. In summary, FABP proteins are vital for normal oxygen consumption, mitochondrial ATP production, and adaption to increased demands for energy, and that their inhibition decreases mitochondrial function.

Having observed decreased cell number by BLI, suppressed metabolic activity, and many changes observed in RNA-Seq and proteomics related to proliferation and cell viability, we next investigated the influence of FABPi on cell cycle and apoptosis in MM.1S and RPMI-8226 cells. The G0/G1 population increased with FABPi combination treatment at 24 hours, which persisted through 72 hours, and a decrease in G2/M was seen at 48 and 72 hours, suggesting a G0/G1 arrest and a negative impact on cell cycle progression in GFP^+^/Luc^+^MM.1S cells(Fig. 5A). Apoptosis was also increased by the combination treatment (Fig. 5B). Apoptosis and cell cycle arrest also occurred in RPMI-8226 cells treated with the combination of inhibitors, or single inhibitors (50 µM or 100 µM) (Supplemental Fig. 17). Interestingly, some cell responses differed after 100 µM combination of inhibitors (50 µM BMS309403 + 50 µM SBFI-26) compared to 100 µM of single inhibitors, suggesting slightly different actions of the inhibitors. Overall, the findings demonstrate that FABPi decrease MM cell numbers by impairing cell cycle progress and inducing apoptosis.

We subsequently investigated the combination of FABPi with dexamethasone, a common first-line therapy for MM patients. FABPi enhanced dexamethasone’s efficacy *in vitro* in GFP^+^/Luc^+^MM.1S, OPM2, and RPMI-8226 (Supplemental Fig. 18A-C). This was in part due to increased apoptosis, with the 3-way combination treatment of BMS309403, SBFI-26, and dexamethasone eliciting the most apoptosis in all three cell lines (Supplemental Fig. 18D-F). In sum, FABPi treatment *in vitro* elicited multitudinous changes in MM cell transcriptome and proteomes signaling alterations in MYC signaling, cellular metabolism, and cell viability which were confirmed by functional analyses.

### FABPi decrease tumor burden and improves survival in xenograft and syngeneic myeloma mouse models

To investigate the efficacy of treating myeloma cells with FABPi *in vivo*, we utilized two murine myeloma models. First, we examined the efficacy of inhibitor treatments in the SCID-beige/ GFP^+^/Luc^+^MM.1S xenograft model. Treatments began with 5 mg/kg BMS309403, 1 mg/kg SBFI-26, the combination, or vehicle 3X/week (Supplemental Fig. 19A) one day after GFP^+^/Luc^+^MM.1S tail vein inoculation. Bone mineral density, but not bone mineral content, was slightly lower after BMS309403 treatment (Supplemental Fig. 19B, C) and fat mass, but not lean mass was decreased with the combination treatment (Supplemental Fig. 19D, E), and FABPi did not influence mouse weight (Fig. 6A). A significant difference in tumor burden assessed by BLI was detected as early as day 21 with all FABPi versus vehicle-treated mice, and this difference continued throughout the study (Fig. 6B, C). Consistent with reduced tumor burden, mice receiving FABPi survived longer than the vehicle-treated mice (Fig. 6D). Similarly, using a second model, the 5TGM1/KaLwRij syngeneic model (Supplemental Fig. 20A), mice treated with 5 mg/kg BMS309403 showed increased survival (Fig. 2E). No adverse effects were observed in either model in response to FABPi, suggesting a good safety profile (Supplemental Fig. 20B).

### Elevated expression of *FABP5* in MM cells corresponds to worse clinical outcomes for patients

To establish potential clinical relevancy, we next tested for an association between FABP5 and MM in independent patient datasets using CoMMpass and OncoMine. In the Multiple Myeloma Research Foundation (MMRF) CoMMpass database, ∼70% of myeloma patient cases exhibited moderate-to-high expression of *FABP5* (defined as >10 counts) (Supplemental Fig. 21A). *FABP3*, *FABP4*, and *FABP6* were expressed by MM cells at lower levels (Supplemental Fig. 21A, insert). We next tested for an association between FABP5 and MM in independent microarray datasets using OncoMine. The Zhan dataset [29] indicated that patients with higher MM cell *FABP5* expression had significantly shorter overall survival (OS) than those with lower expression [30], (Fig. 7A, B), which was confirmed in the Mulligan dataset [31] (Fig. 7C). Similarly, the Carrasco dataset showed a shorter progression-free survival (PFS) in MM patients with high versus low *FABP5* expression (Fig. 7D) [32]. Moreover, patients of the high-risk/poor prognosis subtype had higher *FABP5* expression than those in the more favorable subtypes [30] (Fig. 7E). In the Chng dataset [33], relapsed patients showed increased *FABP5* expression versus newly-diagnosed patients (Fig. 7F). Worse PFS and OS in patients with elevated *FABP5* expression levels was then confirmed in the CoMMpass dataset (log-rank-value for high vs. low expression, <0.0001 for both PFS and OS) (Supplemental Fig. 21B, C). In the Cox proportional hazards model, high *FABP5* expression was associated with a 64% increased risk of disease progression or death (HR: 1.64; CI: 1.34, 2.00), and a 2-fold increased risk of early death (HR: 2.19; CI: 1.66, 2.88).

**Figure 7.**
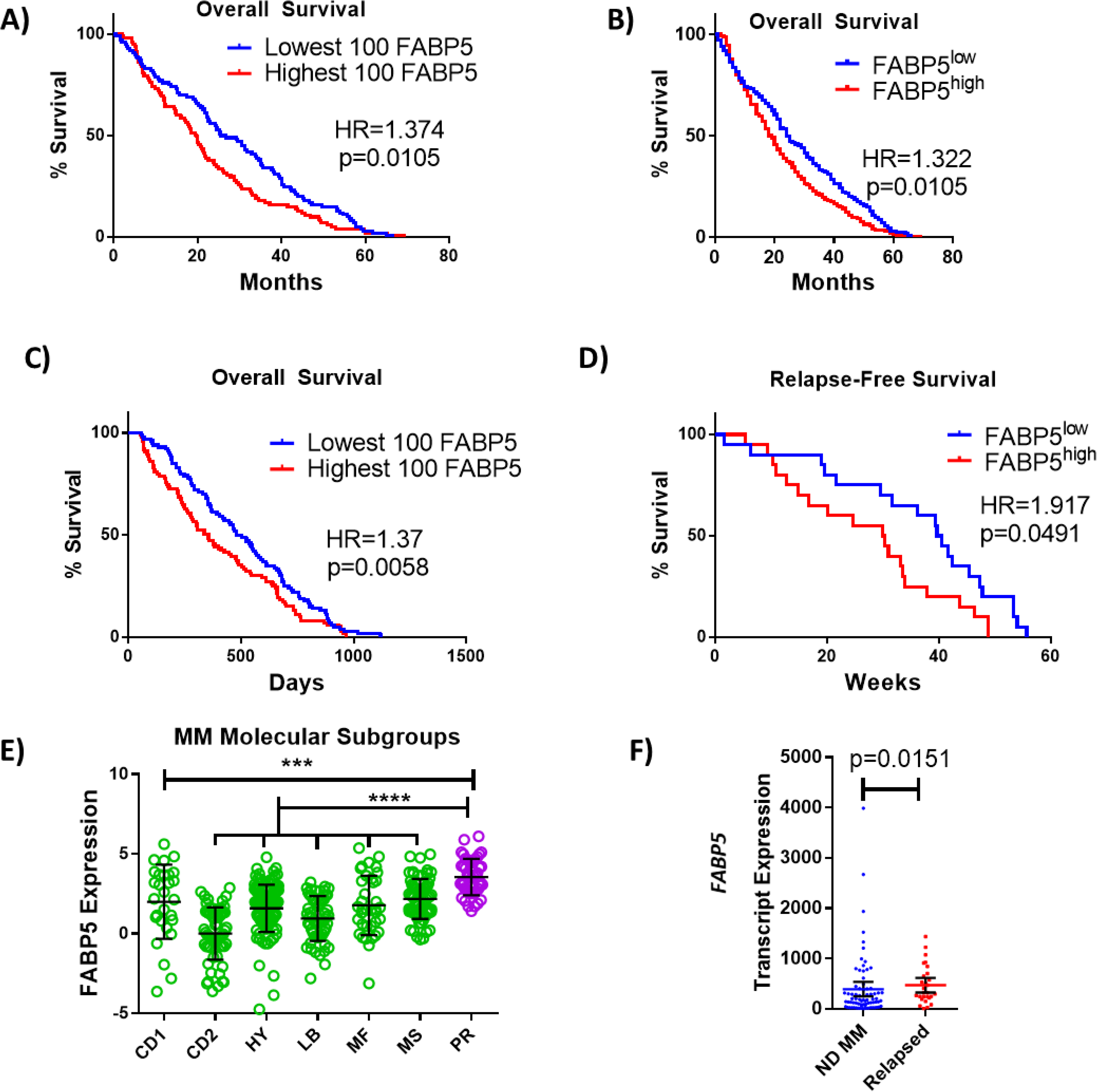
FABP proteins are clinically relevant in MM. A, B) Kaplan-Meier analysis of overall survival (OS) of MM patients in Zhan et al. dataset stratified as top (n=100) or bottom (n=100) *FABP5* expressing, or all patients above (n=207) or below (n=207) the median. C) Kaplan– Meier analysis of relapse-free survival of MM patient groups in Mulligan et al. dataset stratified as top (n=100) or bottom (n=100) *FABP5* expressing. D) Kaplan–Meier analysis of relapse-free survival of MM patient groups in Carrasco et al. dataset: high (n=20) and low (n=20) *FABP5* relative to median. E) Molecular subtypes of MM cells were analyzed for FABP5 expression and significance between all groups and the highly aggressive subtype (PR) was observed using a one-way ANOVA with Dunnett’s multiple comparison testing. (CD1 or CD2 of cyclin D translocation; HY: hyperdiploid; LB: low bone disease; MF or MS with activation of MAF, MAFB, or FGRF3/MMSET; PR: proliferation. From reference [36]). F) Data from Chng et al. dataset from newly-diagnosed (ND) (n=73) and relapsed MM patients (n=28) as mean with 95% confidence interval (CI), with statistical analysis performed using a Mann Whitney test. Data are mean ± SD unless otherwise stated. *p<0.05. **p<0.01. ***p<0.001. ****p<0.0001.

Since obesity is a known MM risk factor [34] and FABP5 can regulate diet-induced obesity [35], we explored the influence of body mass index (BMI) on our findings in the CoMMpass dataset. BMI was not associated with *FABP5* in a general linear model adjusting for age or sex, and the addition of BMI to the Cox model of *FABP5* expression described above did not materially attenuate the effect estimates, suggesting *FABP5* expression is a BMI-independent biomarker for MM aggressiveness. We also examined genes correlated with *FABP5* and found none ontologically related to obesity, again suggesting that FABP5 effects are BMI-independent (Supplemental Fig. 21D; Additional File 1, Supplemental Table 22). All other FABPs expressed in MM cells (*FABP3*, *FABP4*, and *FABP6*) were examined and only *FABP6* showed hazard ratio effects (although effect sizes were not as large as *FABP5*) in PFS (HR:1.48; CI 1.172, 1.869) and OS (HR:1.837, CI: 1.347, 2.504), indicating that FABP6 may also be a biomarker for worse outcomes (Supplemental Fig. 22). Overall, these data across multiple datasets provide rationale to explore the molecular and functional roles of the FABPs in the MM setting.

## DISCUSSION

Herein, we describe our finding that the Fatty Acid Binding Proteins are a novel family of targetable proteins that support myeloma cells. We propose that targeting the FABP family may be a new, efficacious method to inhibit MM progression that necessitates further investigation. FABP inhibition induced apoptosis, cell cycle arrest, and the inhibition of proliferation of numerous MM cell lines *in vitro*, while having negligible effects on non-MM cells. *In vivo* we observed a good safety profile for two different FABP inhibitors, and their combination, supporting prior data demonstrating their safety *in vitro* (in non-cancerous cells) and *in vivo* at doses similar to or above those used here [13,14,17,37]. Myeloma cell proliferation also decreased with genetic knockout of *FABP5,* although FABP signaling compensation may have occurred via upregulation of *FABP6*. Clinical datasets and DepMap analyses also demonstrated the importance of the FABPs, specifically *FABP5,* in MM and implicated this isoform as the most vital for MM cells.

In response to FABPi, we observed decreased expression of genes in XBP1-related and UPR pathways. For example, *EIF5B* was downregulated by all FABPi in proteomic analysis and RNA-Seq. *EIF5B* is a translation initiation factor that promotes the binding of subunits and antagonizes cell cycle arrest via modulations of p21 and p27, and depletion of *EIF5B* could contribute to activation of ER stress [38]. Increased expression of eIF5B has been implicated as a oncoprotein that aids in managing ER stress and evading apoptosis [38]. Myeloma cells constitutively activate the UPR to protect themselves from ER stress-induced death that would otherwise result from the continuous production and secretion of immunoglobulins. Therefore, the inhibition of the protective UPR appears to be one mechanism by which FABPi damage MM cells. We also observed decreased *XBP1* expression and decreased XBP1 pathway activation with FABPi. Based on studies demonstrating the IRE/XBP1 pathway is required for differentiation and survival of MM cells [39], this could be a driver of the decreased UPR and MM cell death resulting from FABPi. Interestingly, decreased UPR and XBP1 signaling could result from decreased MYC expression directly, based on findings that MYC directly controls IRE1 transcription by binding to its promoter and enhancer [40].

While others have shown that BMS309403 reduces UPR in skeletal muscle cells [37], this has not previously been shown in tumor cells before now. As a transcription factor, c-MYC can act as an activator or repressor through either direct binding to regulatory regions, or through chromatin modulation. A MYC activation signature is seen in 67% of MM patients [41], and this signature influences the progression from monoclonal gammopathy of undetermined significance (MGUS) to MM. Targeting MYC in MM cells by knockdown [42] or treatment with a small molecule inhibitor [43] induces cell death; however, the importance of MYC in many healthy cell types make targeting it difficult. Thus, our study represents a novel approach to reducing MYC by targeting the FABP family. This work also builds upon data that myeloma cells exhibit aberrant amino acid, lipid, and energy metabolism [44], and data revealing the importance of metabolic enzymes in myeloma tumorigenesis [45] and drug resistance [46] by demonstrating the role of FABPs in MM cell metabolism and mitochondrial integrity. In sum, we identified a new protein family for therapeutic targeting in myeloma, and demonstrated, for the first time, the great potential for inhibiting it in MM.

## Conclusion

Herein we demonstrated the pivotal role of FABPs in myeloma cell survival *in vitro, in vivo* and clinically. Phamaceutical and genetic inhibition of FABPs result in reduced growth, decreased UPR and Myc signaling, and induced apoptosis. Inhibition *in vivo* significantly increased myeloma bearing mouse survival in immunocompetent and deficient models. Patients that have high FABP5 expression within their myeloma cells result in poor survival and favor subtypes that have poor survival/more aggressive phenotype. Collectively, this data demonstrate preclinically the therapeutic avenue of targeting FABPs in multiple myeloma.

## MATERIALS AND METHODS

### Cell Culture

Human myeloma cell lines GFP^+^/Luc^+^MM.1S (MM.1S), RPMI-8226 (ATCC, Manassas, VA), MM.1R (ATCC), OPM2 (DSMZ), and mouse cell line GFP^+^/Luc^+^ 5TGM1-TK (5TGM1-TK) were maintained in standard MM cell media: RPMI-1640 medium, 10% FBS (Atlanta Biologicals, Flowery Branch, GA), and 1X Antibiotic-Antimycotic (100 U/ml penicillin, 100 μg/ml streptomycin, 0.25 μg/ml fungizone) (ThermoFisher Scientific, Grand Island, NY). U266 (ATCC) cells were maintained in MM growth medium + 15% FBS (Atlanta Biologicals). NCI-H929 (H929, ATCC) cells were maintained in MM growth medium plus 0.05 mM 2-mercaptoethanol. Vk*MYC cells were maintained in RPMI-1640 medium + 20% FBS. Vk*MYC, and MM.1S cells were generously provided by Dr. Ghobrial (Dana-Farber Cancer Institute). GFP^+^/Luc^+^ 5TGM1-TK cells were generously provided by Dr. Roodman (Indiana University). *FABP5* WT and KO MM.1R (ATCC) cells were generated by Synthego (Menlo Park, CA). Primary human MSCs were isolated from deidentified cancellous bone from the acetabulum received from donors (men and women) after total hip arthroplasty through the MaineHealth Biobank after IRB approval and informed consent. Human MSCs were isolated by surface adherence and cultured with a growth media of DMEM, 10% FBS, and 1% an antibiotic-antimycotic as previously described [47–49].

### Cell Number Quantification, Cell Cycle, and Apoptosis *In Vitro* Assays

Cell numbers were measured by bioluminescence imaging (BLI), CellTiter Glo (Promega, Madison, WI), or RealTime Glo (Promega) assays, according to the manufacturer’s instructions, and read on a GLOMAX microplate reader (Promega). Cell cycle analysis was measured with DAPI (0.5 µg/ml) and Ki67 staining (Alexa Fluor 647 Ki67 antibody, 350510, BioLegend). Apoptosis was measured using an annexin V/APC and DAPI Kit (BioLegend); total apoptotic cells were defined as annexin V^+^/DAPI^+^ + annexin V^+^/DAPI^-^ populations. Data were acquired on a Miltenyi MACSquant flow cytometer and data analysis was performed using FlowJo software (BD Life Sciences). For BLI *in vitro* imaging of luciferase expressing cells, sterile luciferin (10µL/well from a 7.5mg/mL stock, VivoGlo, Promega) is added to white, 96 well plates of cells, given 5 minutes to reach equilibrium, and read in a GLOMAX microplate reader (Promega). For flow cytometry, a minimum of 10,000 events was collected and gated off forward and side scatter plots.

### *In Vivo* Experiments

All experimental studies and procedures involving mice were performed in accordance with approved protocols from the Maine Medical Center Research Institute’s (Scarborough, Maine, USA) Institutional Animal Care and Use Committee. In cohort one, eight week old female SCID-beige (CB17.Cg-PrkdcscidLystbg-J/Crl, Charles River) mice were inoculated intravenously (IV) with 5x10^6 GFP^+^/Luc^+^MM.1S cells by a blinded investigator. Treatments then began 3X/week with either 5 mg/kg BMS309403, 1 mg/kg SBFI-26, the combination (5 mg/kg BMS309403 + 1 mg/kg SBFI-26), or the vehicle (5% DMSO), intraperitoneally (n=12/group), based on safe doses reported previously [17, 18]. Body parameters were assessed with piximus at day 1 and 30. In cohort 2, 10-12 week old mice (both sexes, mixed equally between groups) of KaLwRij/C57Bl6 mice (from Dana-Farber Cancer Institute) were injected with 1x10^6 GFP^+^/Luc^+^ 5TGM1-TK cells IV by a blinded investigator, and treated as in cohort one with 5mg/kg BMS309403 (n=9) or vehicle (n=8). Mice were frequently weighed and monitored for clinical signs of treatment-related side effects. “Survival endpoints” were mouse death or euthanasia as required by IACUC, based on body conditioning score including weight loss and impaired hind limb use. Survival differences were analyzed by Kaplan-Meier methodology. For bioluminescent imaging, mice were injected with 150 mg/kg i.p. filter-sterilized D-luciferin substrate (VivoGlo, Promega) and imaged after 15 minutes in an IVIS® Lumina LT (Perkin Elmer, Inc.; Waltham, MA). Data were acquired and analyzed using LivingImage software 4.5.1. (PerkinElmer).

### mRNA Isolation and RNA Sequencing

Three biological sets of GFP^+^/Luc^+^MM.1S cells were cultured for 24 hours with vehicle, 50 μM BMS309403, 50 μM SBFI-26, or the combination prior to mRNA isolation with Qiazol (Qiagen, Germantown, MD) and miRNeasy Mini Kit with on-column DNAse digestion (Qiagen) according to the manufacturer’s protocol. Samples underwent library preparation, sequencing, and analysis at the Vermont Integrative Genomics Resource. See Supplemental Methods for more details.

### Cancer Dependency Map (DepMap) Analysis

Genetic dependency data from the Dependency Map (DepMap) Portal’s CRISPR (Avana) Public20Q3 (https://depmap.org/portal/download/) of 20 human MM cell lines were analyzed and the dependency score (computational correction of copy-number effect in CRISPR-Cas9 essentiality screens (CERES)) of Hallmark Fatty Acid Metabolism genes from Gene Set Enrichment Analysis (https://www.gseamsigdb.org) were determined.

### Survival and Expression Analyses of Clinical Datasets

The Zhan et al. [30] (GSE132604), Carrasco et al. [32] (GSE4452), and Mulligan et al. [31] (GSE9782) datasets were analyzed using OncoMine (ThermoFisher). The Chng dataset [33] showing patient FABP5 mRNA transcript data was analyzed from accession number GEO:GSE6477. The relationship between *FABP5* and MM progression was analyzed with Kaplan-Meier analysis using log-rank Hazard Ratio (HR) and Gehan-Breslow-Wilcoxon significance testing. Gene expression data were downloaded (GEO; GSE6477), log-transformed, and analyzed with an one-way ANOVA model using the aov() function in R, as previously described [50].

For survival analysis in the CoMMpass dataset, survival and Transcripts Per Million (TPM)-normalized gene expression data (IA15 data release) were downloaded from the Multiple Myeloma Research Foundation (MMRF)’s Researcher Gateway (6/16/2021). Patient samples drawn at timepoints other than the baseline were removed from consideration. Based on the histogram of FABP5 expression levels in the CoMMpass cohort, FABP5 expression follows a right-tailed distribution, whereby a subset of patient tumors exhibit higher levels of FABP5. We discretized FABP5 expression based on the cohort’s mean (10.838), stratified samples as FABP5-high and FABP5-low and plotted Kaplan-Meier curves to showcase its effect on OS and PFS. To derive effect estimates, we examined associations between FABP5-high (vs. FABP5-low) in a Cox proportional Hazards Model. Exploratory general linear models also examined the association between BMI and FABP5 expression levels, adjusting for age and sex. Based on the boxplot generated to identify related FABP gene expression levels, FABP3, FABP4 and FABP6 were also significantly expressed in myeloma cells. Thus, following similar procedures, analyses were also conducted based on the cohort’s mean for FABP3 (3.2611), FABP4 (1.624), and FABP6 (0.786).

**Materials and Reagents, Immunofluorescence and Confocal Microscopy, Dual-energy X-ray absorptiometry, CRISPR/Cas9 FABP5-Knockout MM.1R Cell line Development, Western Blotting, Seahorse Metabolic Assay, TMRE Mitochondrial Membrane Potential Assay, qRT-PCR, and Mass Spectrometry Proteomics Methods, Cell Line Validation:** See Supplemental Methods for details.

### Statistical Analysis

Data were analyzed using GraphPad Prism v.6 or above, and unpaired Student’s t tests or one-way or two-way ANOVA using Tukey’s correction was performed, unless otherwise stated. Data are expressed as mean ± standard error of the mean (SEM) or standard deviation (SD); ****p≤ 0.0001; ***p<0.001; **p<0.01; *p<0.05.

### Availability of data and materials

The clinical datasets used and analyzed during the current study are from Oncomine or data related to accession number GEO:GSE6477. RNA-seq data have been deposited in the NCBI Gene Expression Omnibus (GEO) database with the accession number GSE190699. The mass spectrometry proteomic data have been deposited to the ProteomeXchange Consortium via the PRIDE partner respository with the dataset identifier PXD032829.

## AUTHOR CONTRIBUTIONS

M.F. and H.F. designed the experiments, performed *in vitro* and *in vivo* studies, performed pathway and data analysis, and wrote the manuscript. M.K., A.D’A., and C.F performed *in vitro* and *in vivo* experiments. C.S.M. performed DepMap analysis. R.S.P, A.C., C.R.M: performed CoMMpass data analysis. J.A.D. performed RNA-sequencing. L.M. and R.D.I. performed qRT-PCR, E.J. performed flow cytometry. V.D. performed Seahorse metabolic analysis. C.G. and C.V. performed mass spectrometry and proteomic analysis. M.R.R. performed *in vivo* experiments and data analysis, supervised the project, and wrote the manuscript.

## ACKNOWLEDGEMENTS

We thank Dr. Christine Lary for downloading and processing the GSE6477 dataset and Lauren Lever, Samantha Costa, Dr. Clifford Rosen, Madeleine Nowak, Dr. Matt Lynes, and the Vermont Integrative Genomics Resource staff for intellectual or technical contributions. The content is solely the responsibility of the authors and does not necessarily represent the official views of the NIH.

## ADDITIONAL FILES

Additional File 1. Supplemental Tables and Supplemental Table Legends (excel, Additional File 1.xlsx)

## SUPPLEMENTAL INFORMATION

### Supplemental Methods

### Materials and Reagents

Recombinant FABP4 (10009549) and FABP5 (10010364) were purchased from Caymen Chemical (Ann Arbor, MI). Dexamethasone (dex) (VWR), BMS3094013 (Caymen Chemical), SBFI-26 (Aobious, Gloucester, MA), and the MYC inhibitor 10058-F4 (Abcam, Cambridge, UK) were dissolved in DMSO. *In vitro*, dex was used at 80 µM; BMS309403 and SBFI-26 were used at 50 µM either as single treatments or in combination, unless otherwise stated.

### Immunofluorescence and Confocal Microscopy

Patient myeloma cells were fixed and permeabilized using the Nuclear Factor Fixation and Permeabilization Buffer Set (Biolegend, San Diego CA), stained with DAPI (20 µg/ml), antibodies against FABP5 (MA5-2402911215, 1.25 µg/mL, ThermoFisher), and Alexa Fluor 647 anti-rabbit secondary antibody (A-21244, 1.25 µg/mL, ThermoFisher). Cells were then rinsed twice with PBS and imaged on a Leica SP5X laser scanning confocal microscope (Leica Microsystems, Buffalo Grove, IL) with Leica LAS acquisition software, using settings as previously described^1^ using a 20× dry objective on 1.5 mm glass-bottomed dishes (MatTek Corporation, Ashland, MA).

### Dual-energy X-ray absorptiometry

Body parameters (BMD, BMC, Lean Mass, and Fat Mass) were measured with PIXImus duel-energy X-ray densitometer (GE Lunar, Boston, MA, USA). The PIXImus was calibrated daily with a mouse phantom provided by the manufacturer. Mice were anesthetized using 2% isoflurane via a nose cone and placed ventral side down with each limb and tail positioned away from the body. Full-body scans were obtained and DXA data were gathered and processed (Lunar PIXImus 2, version 2.1). BMD and BMC were calculated by extrapolating from a rectangular region of interest (ROI) drawn around one femur of each mouse, using the same ROI for every mouse, and lean and fat mass were also calculated for the entire mouse, exclusive of the head, using Lunar PIXImus 2.1 software default settings.

### CRISPR/Cas9 FABP5-Knockout MM.1R Cell line Development and Characterization

An FABP5-KO pool of MM.1R cells and controls were generated by Synthego using the Guide target ACTTAACATTCTACAGGAGT, Guide sequence ACUUAACAUUCUACAGGAGU and PAM recognition sequence GGG. MM.1R were used as they were found to be the most amenable to CRISPR-Cas9 genetic targeting technology. MM.1R cells were obtained from ATCC by Synthego and confirmed as mycoplasma-negative and free from microbial contamination. Control and KO cell pools were provided to the Reagan lab at passage 4 and passage 5, respectively. Single cell clones were not able to be expanded and thus the pooled sample was used. PCR and sequencing primers used for confirmation were: Fwd: TTTCATATATGTAAAGTGCTGGCTC and Rev:TGATACAGCCTATCATTCTAGAAGCT Wild type and edited cells were thawed and allowed to grow for 1 week prior to seeding (5,000 cells/well; 96-well plate with Real Time Glo (RTG)). Cells from both pools were seeded at ∼1 million cells/T25 for 96 hours prior to harvest for RNA (Qiazol). The expression of FABP family members in both experiments was assessed by qRT-PCR.

### Western Blotting

Protein from cell lysates was extracted using RIPA buffer (Santa Cruz, 24948) or Minute™ Total Protein Extraction Kit (Invent Biotechnology, SD-001/SN-002) and quantified using a DC protein assay kit II (Bio-Rad, 5000112). Samples were denatured in 4x laemmli buffer (Bio-Rad, 1610747) with β-mercaptoethanol (VWR, 97064-880) for 5 minutes at 95°C, run on 12% polyacrylamide gels (Bio-Rad, 5671043), and transferred onto PVDF membranes (Bio-Rad, 1704156). Blots were blocked for 2 hours in 5% non-fat milk (VWR, 10128-602). Staining protocols with antibody details are in Supplemental Table 23 (Additional File 1). All antibodies were incubated at 4°C. Blots were imaged after adding ECL reagents (Biorad, 1705060) for 5 minutes and visualized using Azure c600 (Azure biosystems).

### Seahorse Metabolic Assays

In the Maine Medical Center Research Institute’s Physiology Core, MM.1S cells were cultured for 24 hours with BMS309403 (50 µM), SBFI-26 (50 µM), or both and then adhered to Cell Tak (Corning)-coated Seahorse XF96 V3 PS cell culture microplates (Agilent, # 101085-004) at a density of 60,000 cells/well in XF DMEM medium pH, 7.4 (Aglient # 103576-100) supplemented with 1mM sodium pyruvate, 2mM glutamine and 10mM glucose according to the manufacturer’s instructions (https://www.agilent.com/cs/library/technicaloverviews/public/5991-7153EN.pdf).

Oxygen consumption rate in cells was then measured in basal conditions and in response to oligomycin (1.25 µM), FCCP (1 µM), and rotenone and antimycin A (0.5 µM). Data were analyzed using Wave Software V2.6 and Seahorse XF Cell Mito Stress Test Report Generators (www.agilent.com). A one-way ANOVA was used for each parameter with Uncorrected Fisher’s LSD multiple comparison post-hoc testing for significance. Results represent 5 independent experiments with 1 representative experiment shown with 20-24 wells per condition.

In a separate set of experiments, repeated 2 times, cells were treated as above, however etomoxir or vehicle was added at a final concentration of 4 μM prior to subjecting the cells to the mitochondrial stress test. Due to artificial increases in OCR caused by further warming of the plate during ETOX measurements, the ETOX response data was normalized to MM.1S (vehicle, vehicle) control cells.

### TMRE Mitochondrial Membrane Potential Assay

MM.1S cells were cultured for 24, 48 and 72 hours with BMS309403 (50 µM), SBFI-26 (50 µM), or combination before staining with 0.5 mM TMRE for 30 minutes per Caymen Chemical protocol. Data acquisition was performed on a Miltenyi MACSquant flow cytometer and data analysis was performed using FlowJo analysis software (BD Life Sciences) with a minimum of 10,000 events collected and gated off forward and side scatter plots.

### mRNA Isolation and RNA Sequencing Continued

mRNA was quantified and tested for quality and contamination using a Nanodrop (Thermo Fisher Scientific) and subjected to quality control standards of 260/230>2 and 260/280>1.8 prior to library preparation. Partek^®^ Flow (version 10.0.21.0302) was used to analyze the sequence reads. Poorer quality bases from the 3’ end were trimmed (phred score <20), and the trimmed reads (ave. quality > 36.7, ave. length 75 bp, ave. GC ∼56%) were aligned to the human reference genome hg38 using the STAR 2.6 aligner. Aligned reads were then quantified using an Expectation-Maximization model, and translated to genes. Genes that had fewer than 30 counts were then filtered, retaining 14,089 high count genes. Differentially expression comparisons were performed using DESeq2. Downstream comparisons of IPA canonical pathways and upstream regulators were executed in Excel (Microsoft, Redmond, WA). Data were analyzed through the use of IPA^2^ (QIAGEN, https://www.qiagenbioinformatics.com/products/ingenuitypathway-analysis) and STRING DB version 11.0. RNAseq heatmap of Myc pathway was generated on http://www.heatmapper.ca/expression using single linkage and Pearson distance measurement algorithms.

### Quantification of Global 5-Hydroxymethylcytosine Levels

DNA was isolated from 1 million MM.1S cells after 24 hours of treatment with vehicle (DMSO) or 50 µM BMS309403 and 50 µM SBFI-26 using the DNeasy Blood and Tissue kit (Qiagen, Germantown, MD, USA) per the manufacturers instructions. DNA was quantified and tested for quality and contamination using a Nanodrop 2000 (Thermo Fisher Scientific) and subjected to quality control minimum standards of 260/230>2 and 260/280>1.8 prior to use in subsequent steps. 100 ng of DNA was then analyzed via MethylFlash Global DNA Hydroxymethylation (5-hmC) ELISA Easy Kit (Cat.# P-1032-48, Epigentek, Farmingdale, NY, USA) per the manufacturer’s instructions.

### Quantitative RT-PCR

MM.1S, 5TGM1-TK and RPMI-8226 cells were cultured for 24 hours with treatments prior to mRNA isolation as described above. cDNA synthesis (Applied Biosciences High Capacity cDNA Kit, ThermoScientific, Waltham, MA, USA) was executed prior to quantitative PCR (qRT-PCR) using SYBR Master Mix (Bio-Rad, Hercules, CA, USA) and thermocycling reactions were completed using a CFX-96 (Bio-Rad Laboratories). Data were analyzed using Bio-Rad CFX Manager 3.1 and Excel (Microsoft Corp., Redmond, WA, USA) using the delta-delta CT method. Primer details are in Supplemental Table 24 (Additional File 1).

### Mass Spectrometry Proteomics Sample Preparation

1. Cells for proteomics analysis were harvested by scraping into centrifuge tubes and pelleting for 5 minutes at 2,500 × g, 4°C. Cells were then resuspended in PBS and pelleted, twice for a total of two cell pellet washes.
2. Cells were solubilized in ice-cold RIPA buffer and DNA sheared using a probe-tip sonicator (3 × 10 seconds) operating at 50% power with the samples on ice. Each was then centrifuged (14,000 × g) at 4°C and the supernatant collected. Protein content was measured relative to bovine serum albumin protein concentration standards using the bicinchoninic acid (BCA) assay (Thermo Scientific Pierce, Waltham, MA).
3. Approximately 100 µg protein from each sample was used in further sample preparation. Protein precipitation was initiated with the addition of a 10-fold volumetric excess of ice-cold ethanol. Samples were then placed in an aluminum block at -20°C for one hour, then protein pelleted in a refrigerated tabletop centrifuge (4°C) for 20 minutes at 16,000 × g. The overlay was removed and discarded. Protein samples were allowed to dry under ambient conditions.
4. Each sample was resuspended in 50 mM Tris (pH = 8.0) containing 8.0 M urea and 10 mM TCEP (tris(2-carboxyethyl)phosphine hydrochloride, Strem Chemicals, Newburyport, MA). Reduction of cysteine residues was performed in an aluminum heating block at 55°C for 1 hour.
5. After cooling to room temperature, each sample was brought to 25 mM iodoacetamide (Thermo Scientific Pierce, Waltham, MA) and cysteine alkylation allowed to proceed for 30 minutes in the dark. Reactions were quenched with the addition of 1-2 µL 2-mercaptoethanol (Thermo Scientific, Waltham, MA) to each sample.
6. Each was diluted with 50 mM Tris buffer (pH = 8.0 - 8.5) containing 1.0 mM calcium chloride (Sigma-Aldrich, St. Louis MO) such that the urea concentration was brought below 1.0 M. Sequencing-grade modified trypsin (Promega, Madison, WI) was added to a final proportion of 2% by mass relative to sample total protein as measured with the BCA assay. Proteolysis was performed overnight at 37°C in the dark.
7. Samples were evaporated to dryness using a centrifugal vacuum concentrator. Each was redissolved in 4% acetonitrile solution containing 5% formic acid (Optima grade, Fisher Scientific, Waltham, MA). Peptides were freed of salts and buffers using Top Tip Micro-spin columns packed with C18 media (Glygen Corporation, Columbia, MD) according to manufacturer-suggested protocol.
8. Samples were again evaporated to dryness using a centrifugal vacuum concentrator and peptides redissolved in 4% acetonitrile solution containing 5% formic acid (Optima grade).

### LC-MS/MS

All sample separations performed in tandem with mass spectrometric analysis are performed on an Eksigent NanoLC 425 nano-UPLC System (Sciex, Framingham, MA) in direct-injection mode with a 3 µL sample loop. Fractionation is performed on a reverse-phase nano HPLC column (Acclaim PepMap 100 C18, 75 µm × 150 mm, 3 µm particle, 120 Å pore) held at 45°C with a flow rate of 350 nL/min. Solvents are blended from LC-MS-grade water and acetonitrile (Honeywell Burdick & Jackson, Muskegon, MI). Mobile phase A is 2% acetonitrile solution, while mobile phase B is 99.9% acetonitrile. Both contain 0.1% formic acid (Optima grade, Fisher Chemical, Waltham, MA). Approximately 1 µg of peptides are applied to the column equilibrated at 3% B and loading continued for 12 minutes. The sample loop is then taken out of the flow path and the column washed for 30 seconds at starting conditions. A gradient to 35% B is executed at constant flow rate over 90 minutes followed by a 3-minute gradient to 90% B. The column is washed for 5 minutes under these conditions before being returned to starting conditions over 2 minutes.

Analysis is performed in positive mode on a TripleTOF 6600 quadrupole time-of-flight (QTOF) mass spectrometer (Sciex, Framingham, MA). The column eluate is directed to a silica capillary emitter (SilicaTip, 20 µm ID, 10 µm tip ID, New Objective, Littleton, MA) maintained at 2400-2600 V. Nitrogen nebulizer gas is held at 4-6 psi, with the curtain gas at 21-25 psi. The source is kept at 150°C.

Data acquisition performed by information-dependent analysis (IDA) is executed under the following conditions: a parent ion scan is acquired over a range of 400-1500 mass units using a 200 msec accumulation time. This is followed by MS/MS scans of the 50 most-intense ions detected in the parent scan over ranges from 100-1500 mass units. These ions must also meet criteria of a 2^+^-5^+^ charge state and of having intensities greater than a 350 counts-per-second (cps) threshold to be selected for MS/MS. Accumulation times for the MS/MS scans are 15 msec. Rolling collision energies are used according to the equation recommended by the manufacturer. Collision energy spread is not used. After an ion is detected and fragmented, its mass is excluded from subsequent analysis for 15 seconds.

SWATH analysis is performed according to previously-published optimized conditions tailored to the 6600 instrument^3^. Briefly, SWATH MS/MS windows of variable sizes are generated using Sciex-provided calculators. Rolling collision energies are used, as well as fragmentation conditions optimized for ions of a 2^+^ charge state. SWATH detection parameters are set to a mass range of m/z = 100-1500 with accumulation times of 25 msec in the high-sensitivity mode. A parent-ion scan is acquired over a range of 400-1500 mass units using a 250 msec accumulation time. The Pride database was used to upload data^4^, and InteractiveVenn software was used to make Venn Diagrams to combine Proteomic and RNAseq data (http://www.interactivenn.net/#)^5^. Proteomic Heatmaps were generated using centroid linkage and Kendall’s Tau distance measurement algorithms with http://www.heatmapper.ca/expression.

### Cell Line Validation

Data and methods previously described validated the MM.1S, OPM-2, and RPMI-8226^1^. 5TGM1-TK and and Vk*Myc cells have not been validated at this time. Cells were validated as mycoplasma and virus negative by the Yale Comparative Pathology Research Core on the following dates:

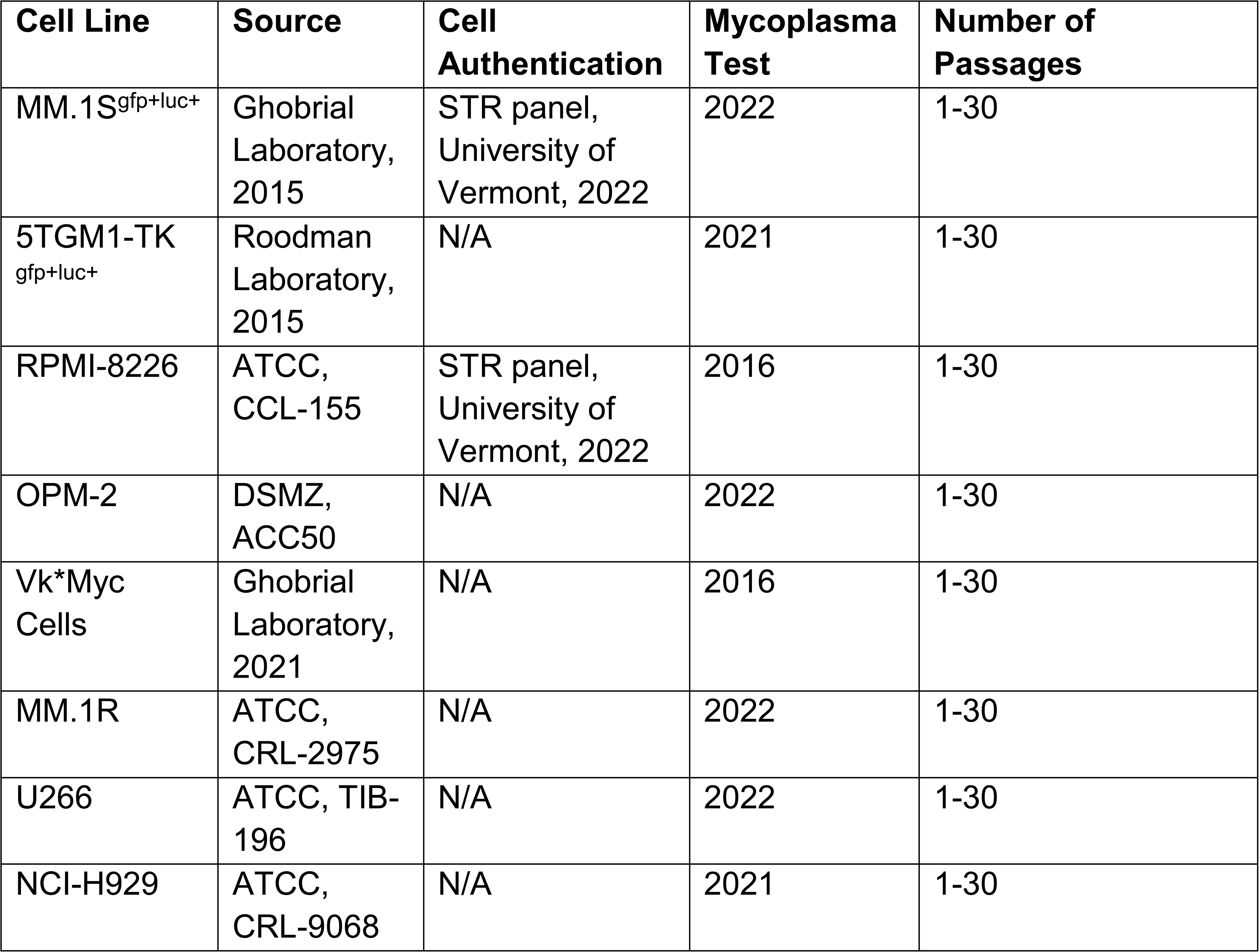

## Supplemental Figures & Legends

**Supplemental Figure 1.**
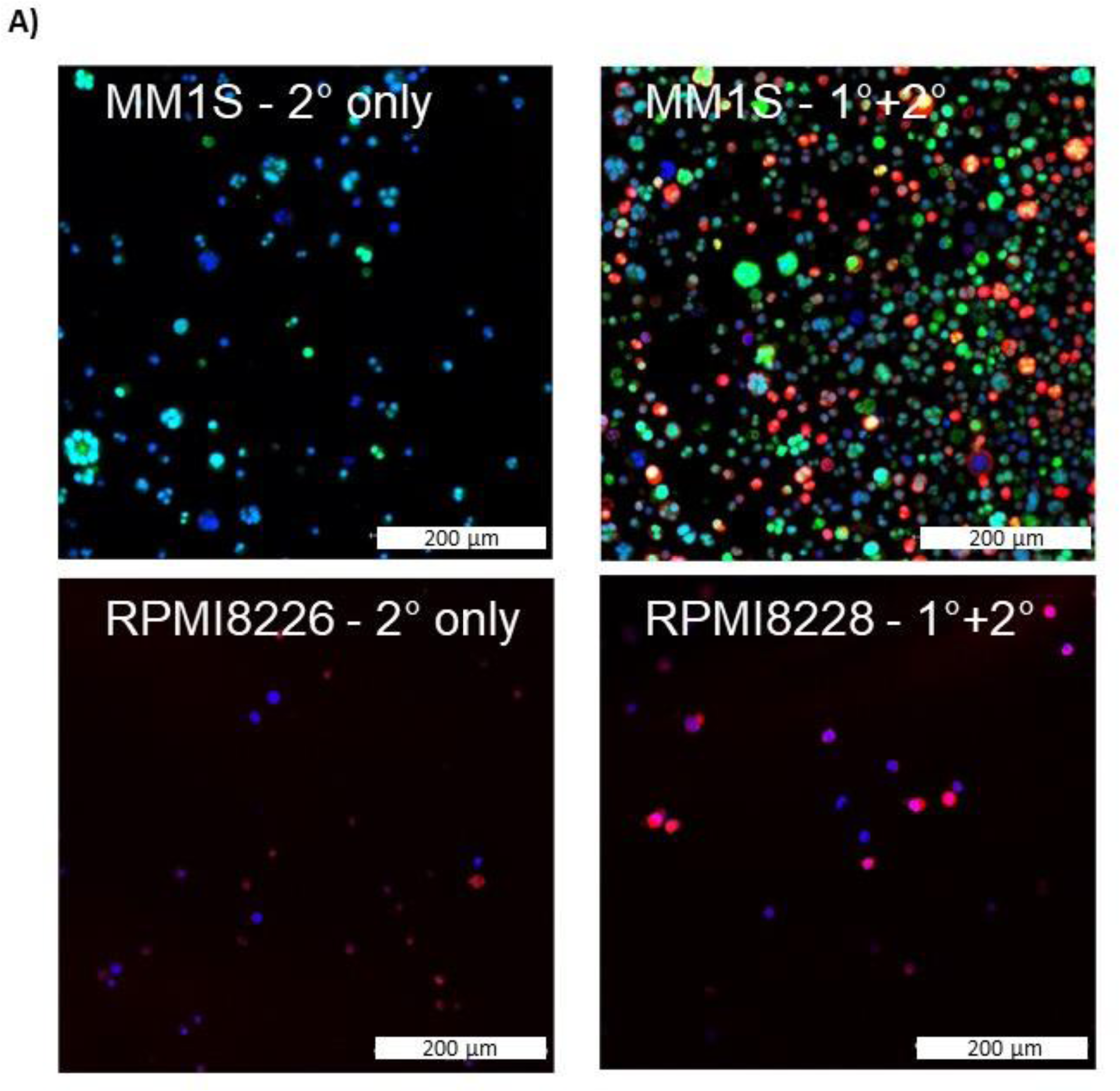
Confocal overlay immunofluorescence images show FABP5 (red) primarily in cytoplasm. Nuclei identified with DAPI (blue) in RPMI8226 and GFP^+^/Luc^+^ MM.1S (green) cells. Cells stained with secondary antibody alone (left) and primary and secondary (right). Scale bar=200 µm.

**Supplemental Figure 2.**
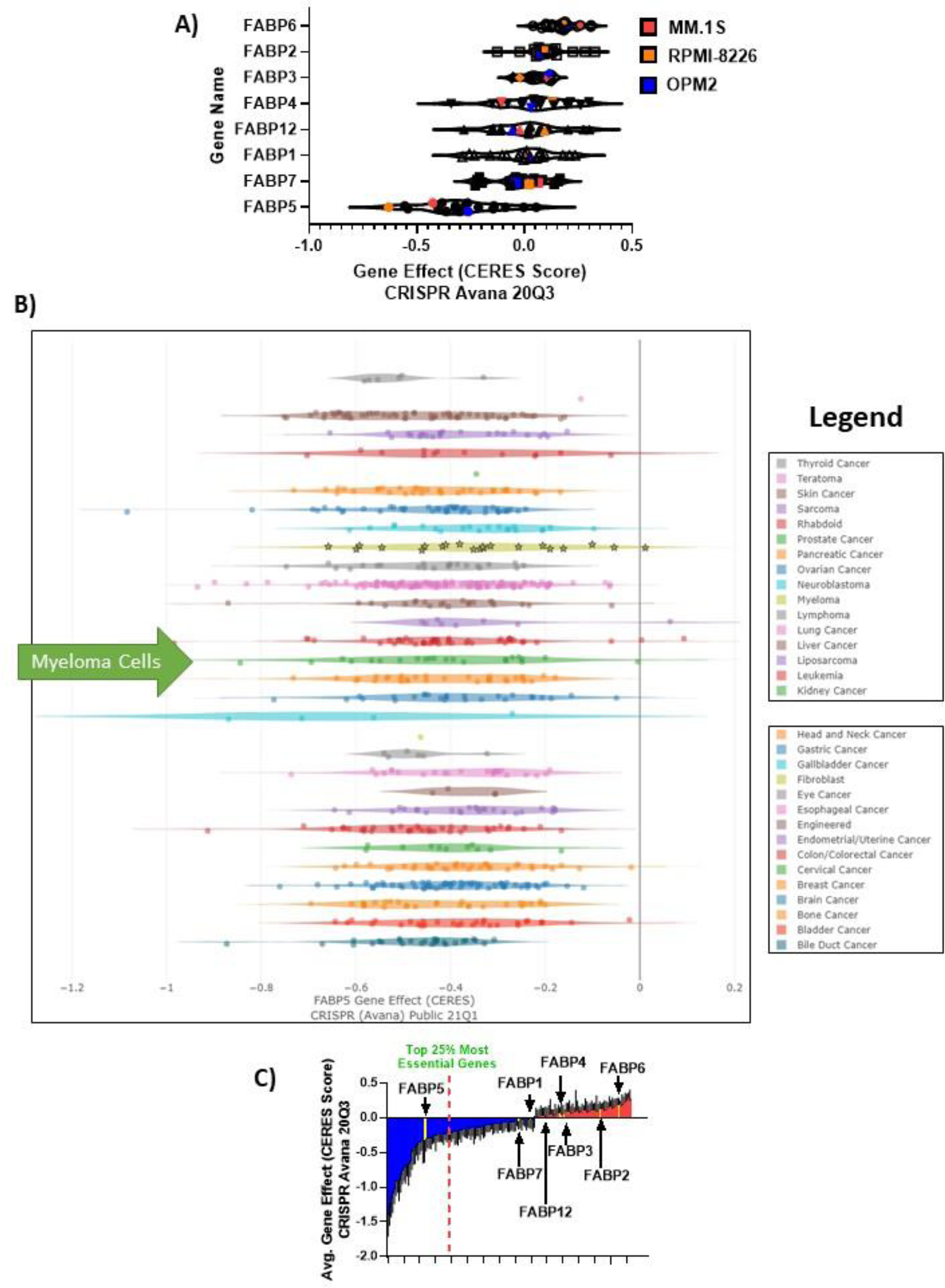
FABP5 Depmap CERES scores in all tumor cells. FABP proteins are clinically relevant in MM. A) CERES DepMap scores for the FABP members labeled with mean scores. B) Derived from CRISPR Avana Public 21Q1 screen for all tumor cells available in Depmap database. FABP5 showed a negative value for all cancer types, demonstrating a dependency on FABP5 for tumor cell survival. Myeloma cells are highlighted with stars and green arrow. Legend provided is in the same order as the cell lines plotted. C) Fatty acid metabolism-related genes in the DepMap; genes with negative scores in blue (essential) and positive scores (less essential) in red.

**Supplemental Figure 3.**
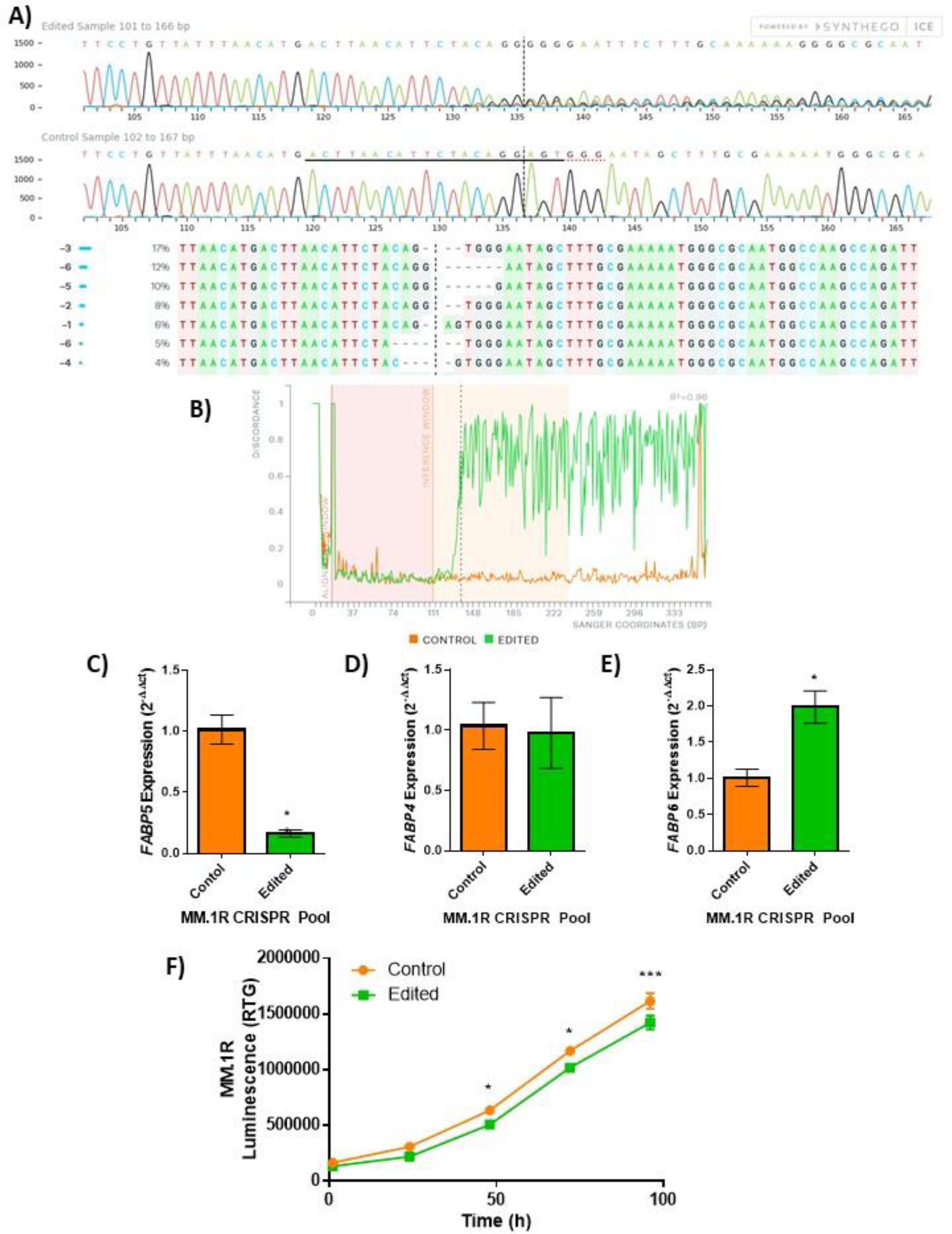
Knockout of *FABP5* in MM.1R myeloma cell line. *FABP5* was genetically targeted by CRISPR-Cas9 in the MM.1R human myeloma cell line (ATCC). A) Sanger sequencing confirmation of mutant (edited) and control (wild type) MM.1R populations including traces (top), and inferred sequences present in MM.1R edited population (middle, wild-type sequence “+”, panel. B) Representation of alignment between the wild type (control, orange) and mutant (edited, green) cells. In panels A and B, dotted lines indicate CRISPR-Cas9 cut site; data provided by Synthego. qRT-PCR analysis of the expression of *FABP* family members in the control (orange) and edited (green) pools: C) *FABP5*, D) *FABP4*, and E) *FABP6*; n=3, data plotted as Mean ± SEM with significance determined by Student’s t-test (*p<0.05). F) RealTime-Glo analysis of MM.1R cells in *FABP5* control (orange) and edited (green) lines over time (n=8 wells per group), significance was determined by 2-way ANOVA with Sidak’s multiple comparisons test (*p<0.05, ***p<0.001). Data plotted as Mean ± SEM for 1 experiment with 8 technical wells; representative of two separate experiments.

**Supplemental Figure 4.**
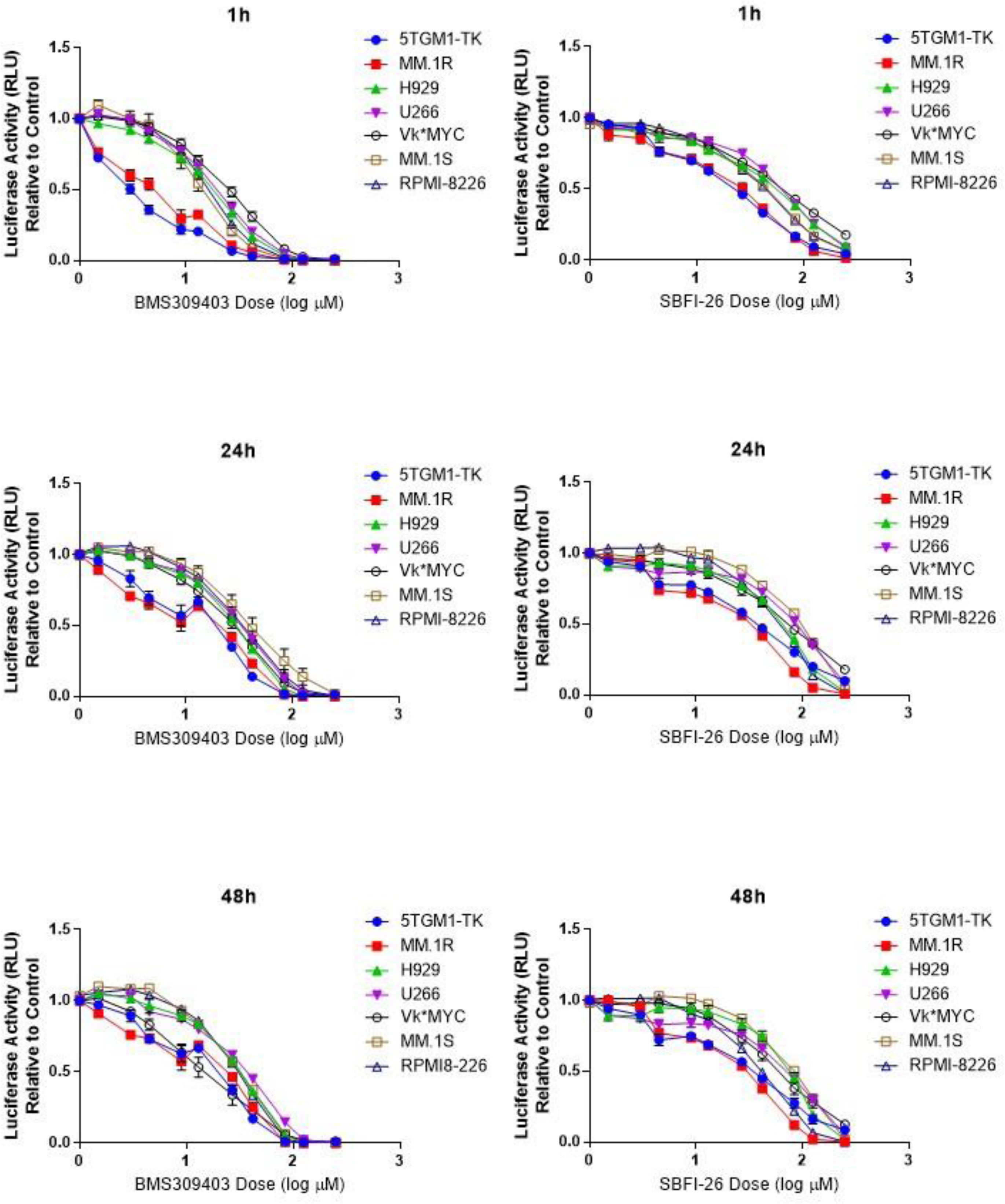
FABP inhibitors exhibit consistent negative effects on cell number in 8 myeloma cell lines. RealTime-Glo analysis of MM cell lines over time with FABP inhibitors demonstrates dose-dependent decreases in luciferase activity. Data represent mean ± SEM from at least 3 biological repeats.

**Supplemental Figure 5.**
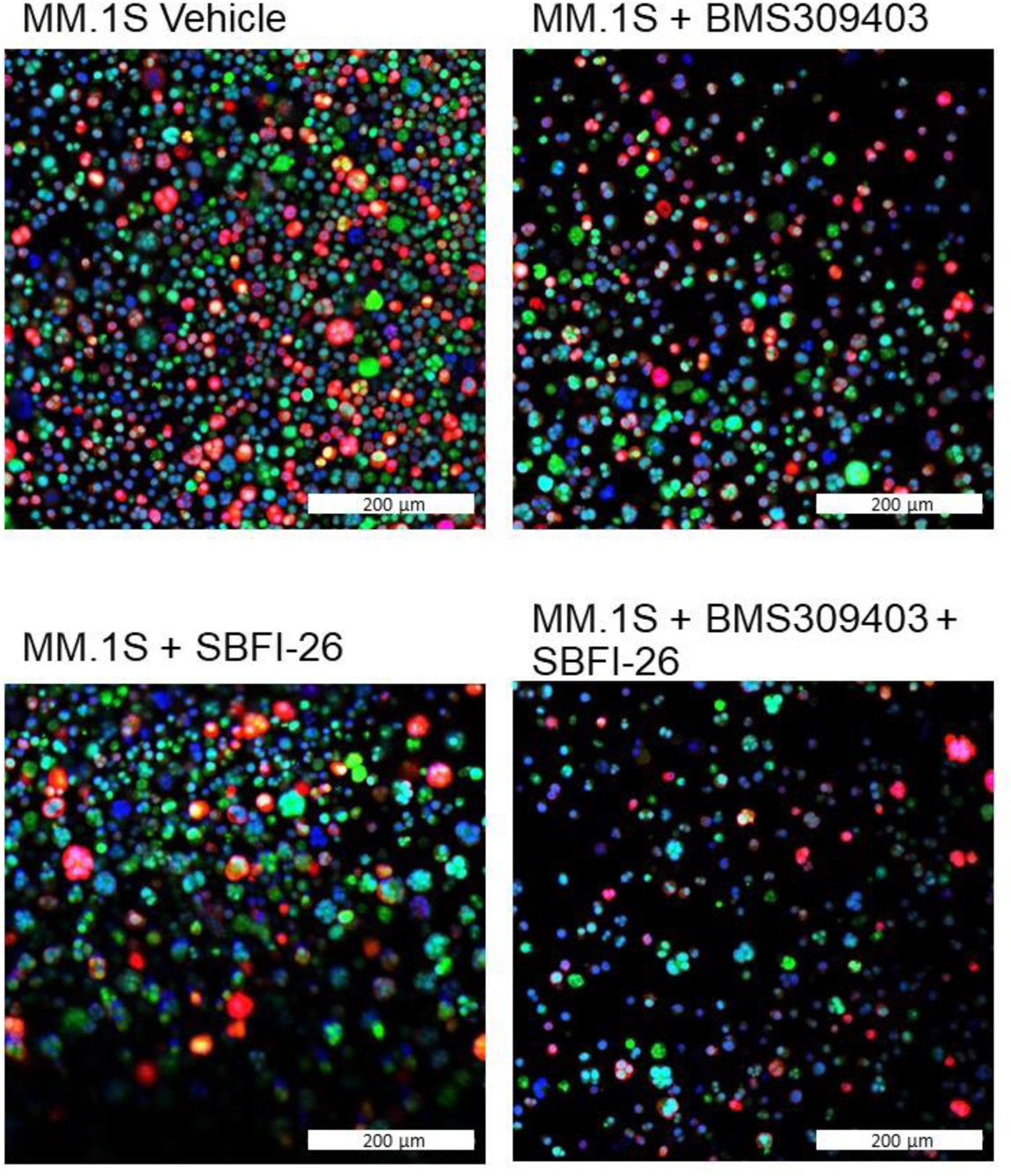
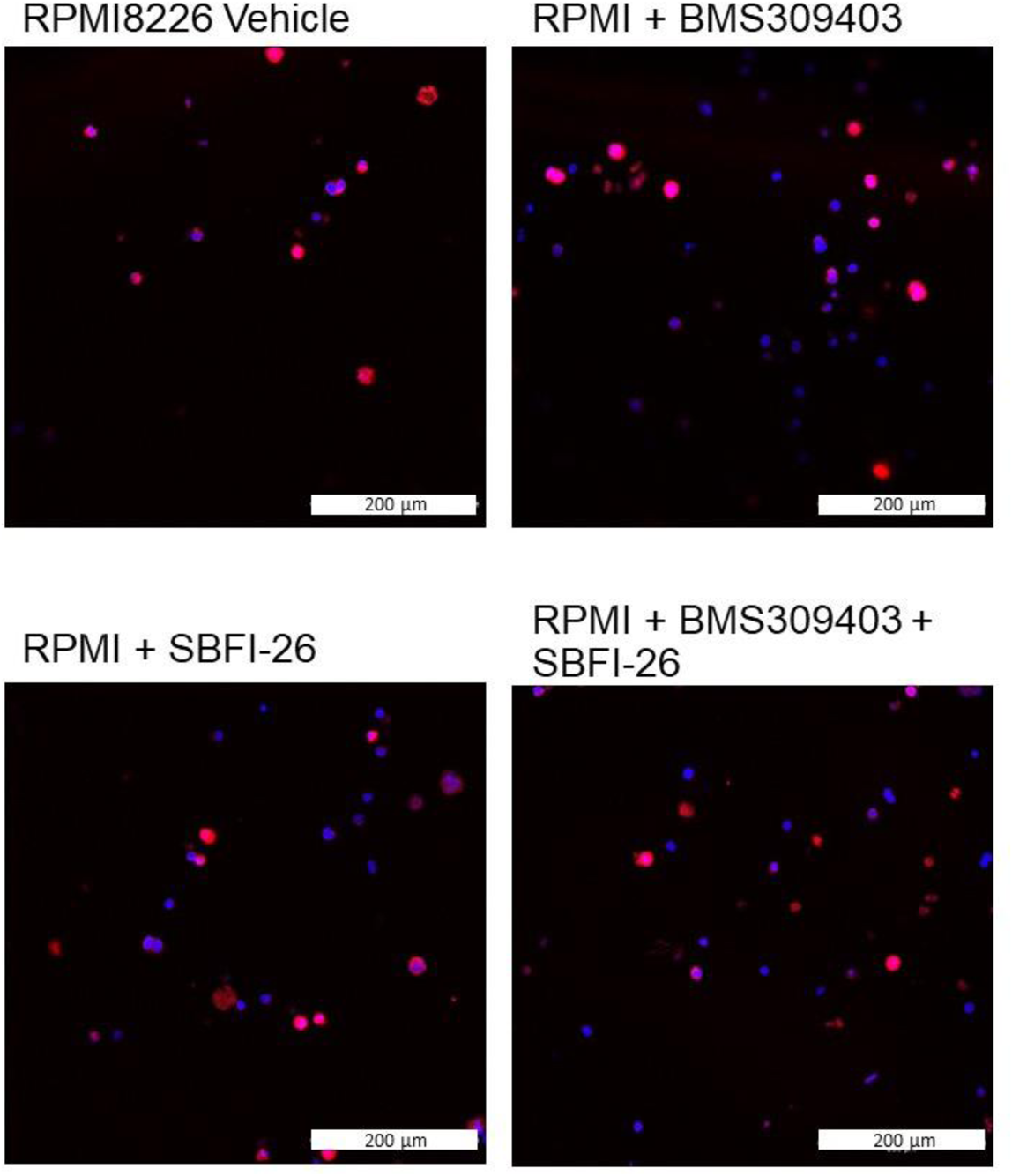
FABP inhibitor treatment did not induce changes in amount or localization of FABP5. Immunofluorescence images with confocal overlay show cells stained for nuclei (DAPI, blue) and FABP5 (red) after treatment with vehicle control or FABP inhibitors (50 µM) for 24 hours. Inhibitors did not appear to alter the expression or location of FABP5 in MM.1S (A) or RPMI-8226 (B) myeloma cells. Representative confocal images from 3 wells.

**Supplemental Figure 6.**
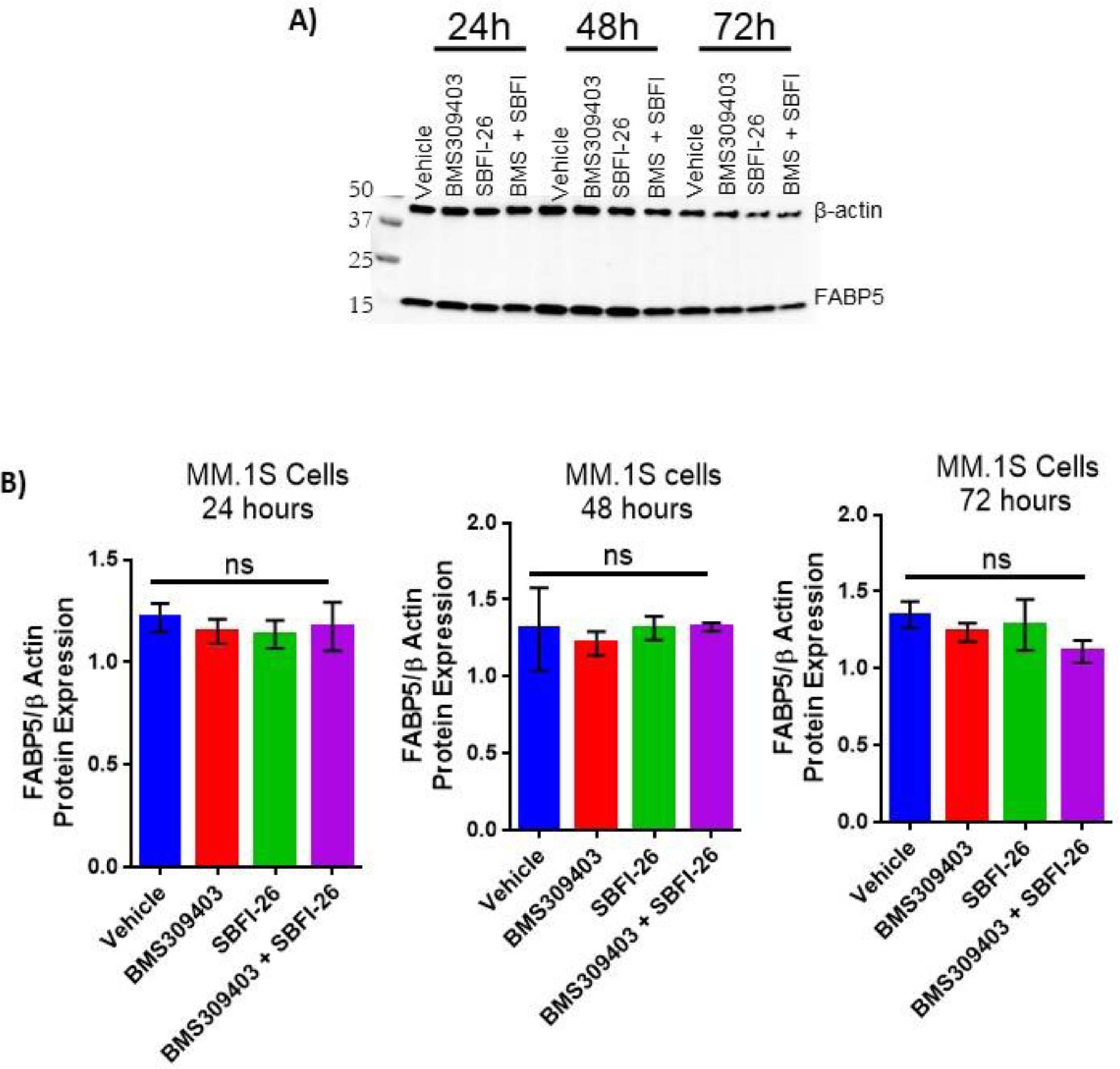
Protein levels of FABP5 not affected by inhibitors in MM.1S cells. A) Representative western blot of B-actin (housekeeping control) and FABP5 protein levels at 24, 48, and 72 hours after treatment with BMS309403 (50uM), SBFI-26 (50uM) or the combination. B) Quantification of Western blots. Data represent Mean ± SEM from 3 biological repeats, analyzed with one-way ANOVA.

**Supplemental Figure 7.**
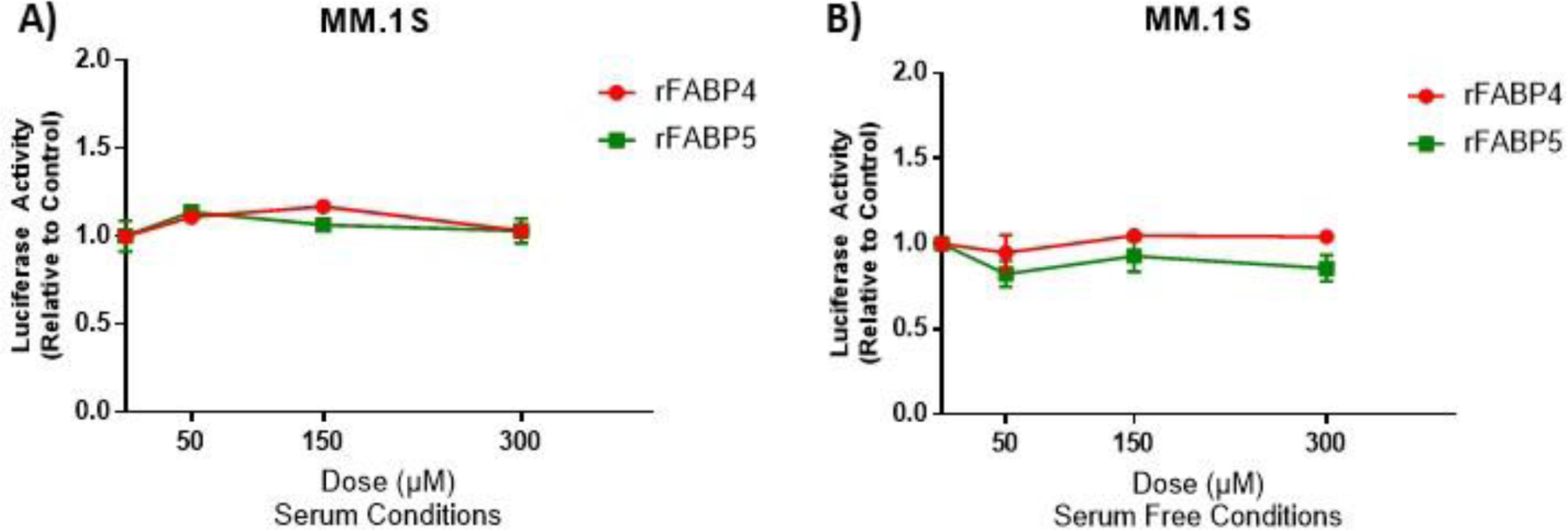
Treatment with Recombinant FABP4 or FABP5 has no effect on cell number in myeloma cell lines. Luciferase activity analysis after 72 hours of recombinant human FABP4 or FABP5 protein treatment in myeloma cells in 10% serum or serum free conditions. A) GFP^+^/Luc^+^MM.1S cell growth was monitored with exogenous luciferin after exposure to recombinant FABP4 or 5 (rFABP4 or rFABP5) protein in serum containing or (B) serum free conditions. Data represent n=3, Mean ± SEM, averages of at least 3 experimental repeats. Statistical analysis was determined with a one-way ANOVA for each recombinant FABP; no significance was observed.

**Supplemental Figure 8.**
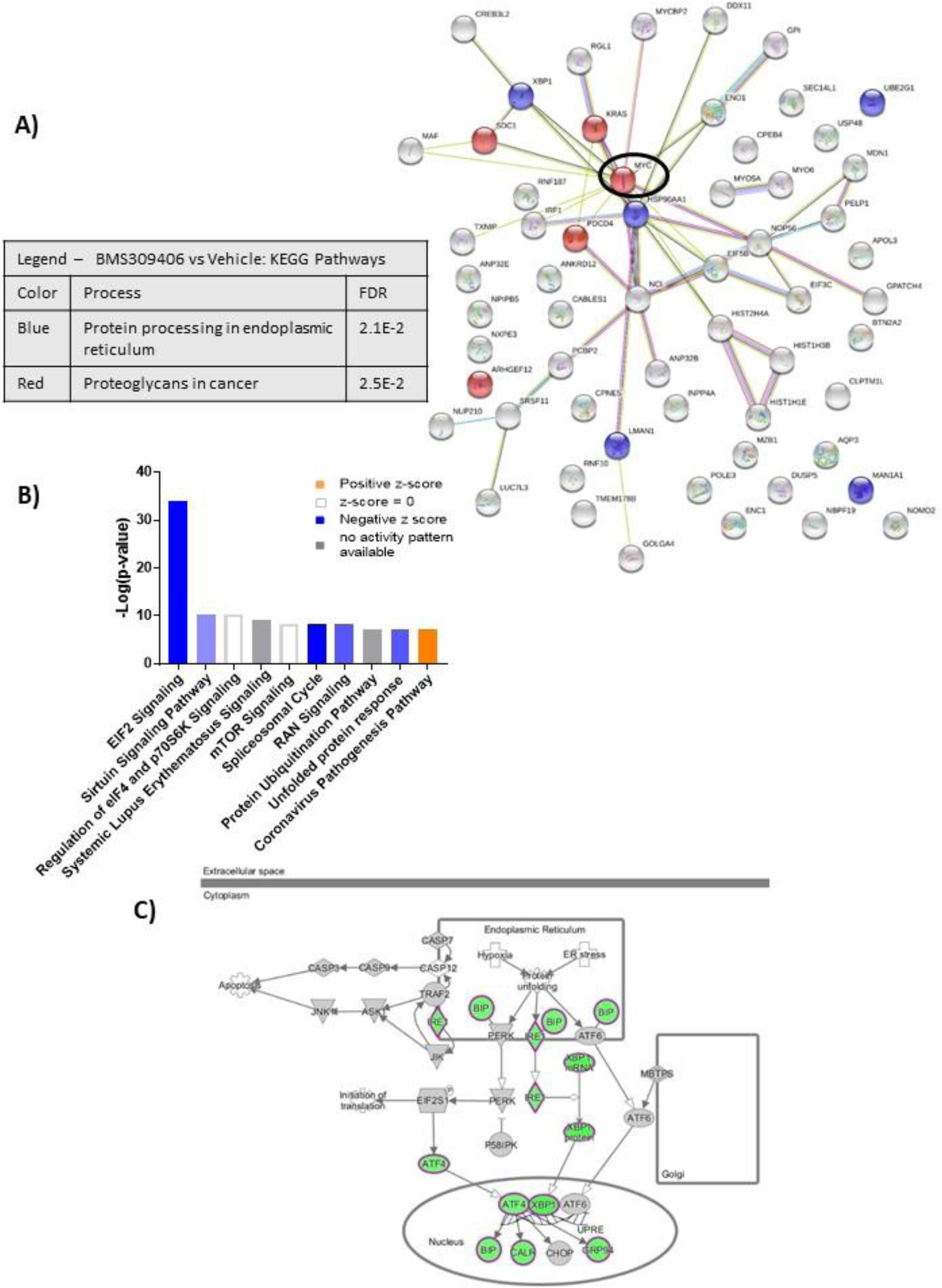
RNA sequencing analysis of GFP^+^/Luc^+^MM1S cells treated for 24 hours with BMS309403. A) String db visualization of significantly altered genes (FDR<0.2) demonstrates importance of MYC as central, connected node (circled). B) Ingenuity pathway analysis (IPA) reveals altered canonical pathways after BMS309403 treatment compared to control such as endoplasmic reticulum stress pathway (C). IPA canonical pathway analysis utilizes p-value of overlap by Fisher’s exact test, significance threshold value of p<0.05(-log value of 1.3).

**Supplemental Figure 9.**
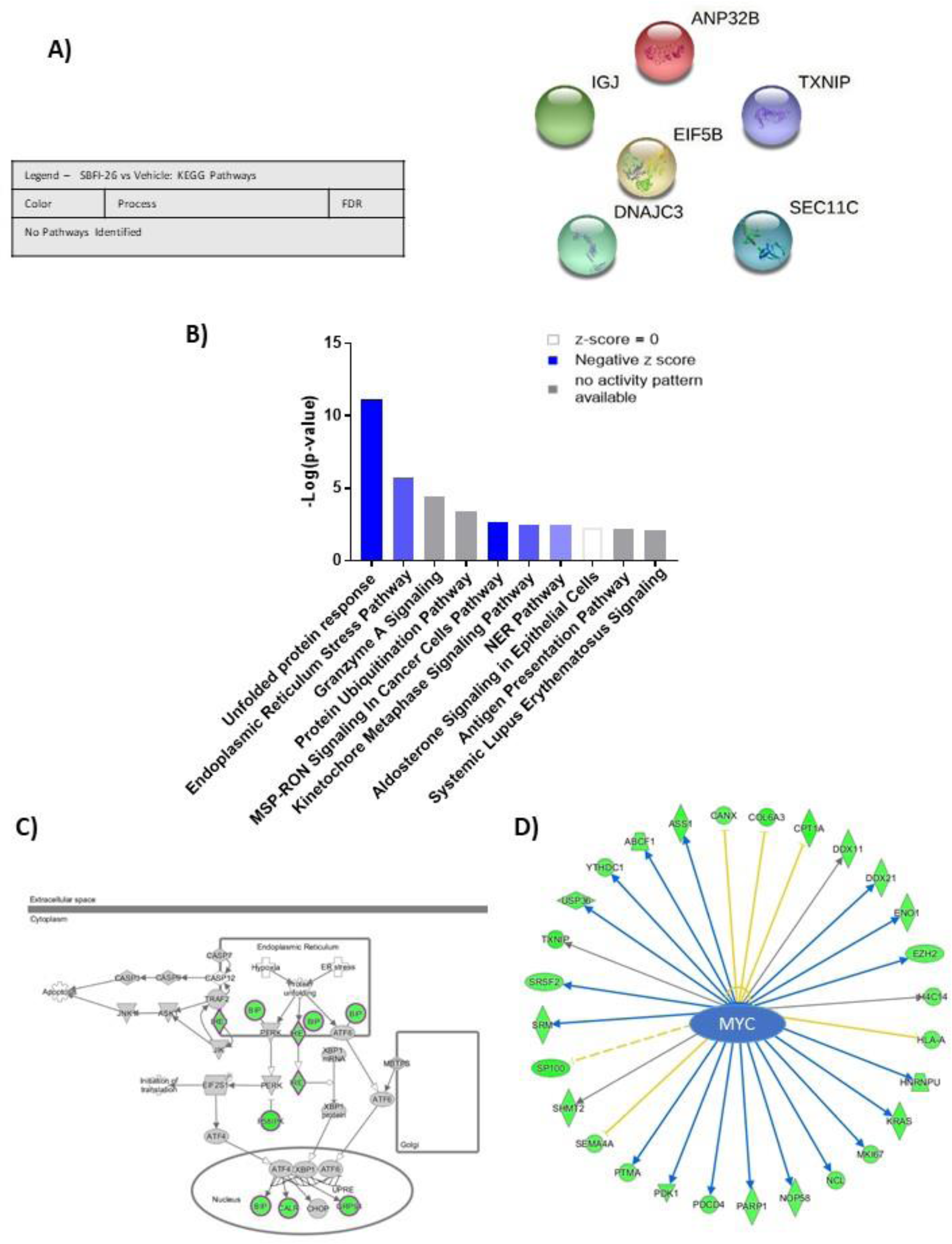
RNA sequencing analysis of MM1S cells treated for 24 hours with SBFI-26. A) Ingenuity pathway analysis prediction show a reduction in canonical signaling pathways after SFBI-26 treatment compared to control such as B) endoplasmic reticulum stress pathway and C) MYC-regulated molecules (arrows indicate expression consistency with predicted patterns (blue=consistent, yellow=unknown, orange=inconsistent; color of molecules indicates expression pattern (green=decreased by treatment, red=increased by treatment)). IPA canonical pathway analysis utilizes p-value of overlap by Fisher’s exact test, significance threshold value of p<0.05(-log value of 1.3).

**Supplemental Figure 10.**
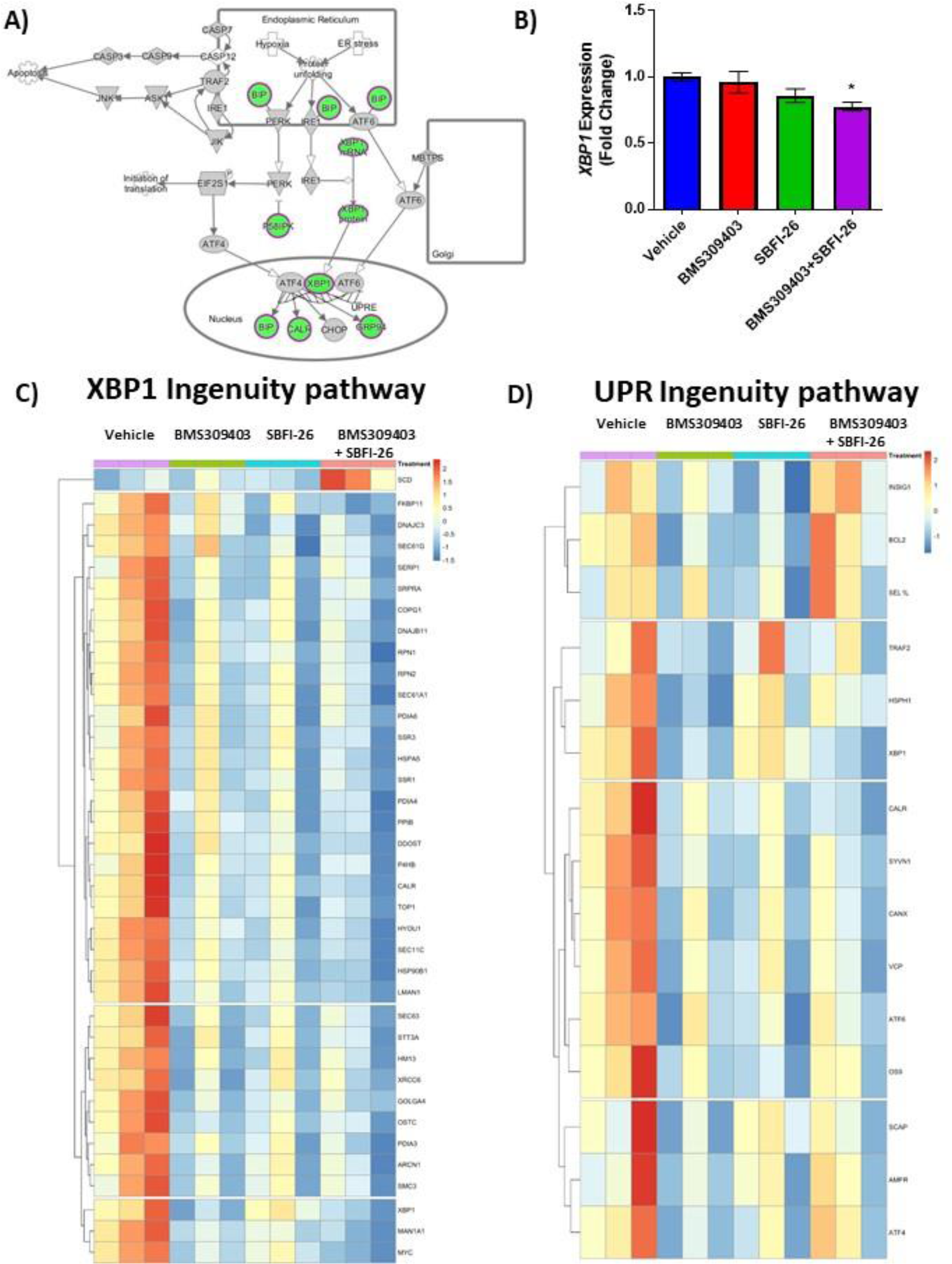
RNA sequencing analysis of MM1S cells treated with FABP inhibitors reveals a unique gene expression suggesting endoplasmic reticulum stress. A) IPA ER Stress for Co-treatment vs. Vehicle, where green is decreased and grey is not changed. B) Semi-quantitative RT-PCR assessment of total *XBP1* transcripts in MM.1S cells after 24 hour treatments with BMS309403 (50 µM), SBFI-26 (50 µM) or the combination normalized to RPLP0 housekeeping gene. Data represent mean ± SEM from n=3 biological repeats, analyzed with one-way ANOVA with Dunnett’s multiple comparisons test; significance shown as *p<0.05. C) Heatmap visualization of genes involved in XBP1 signaling (D) and the unfolded protein response (UPR) as determined by Ingenuity Pathway Analysis.

**Supplemental Figure 11.**
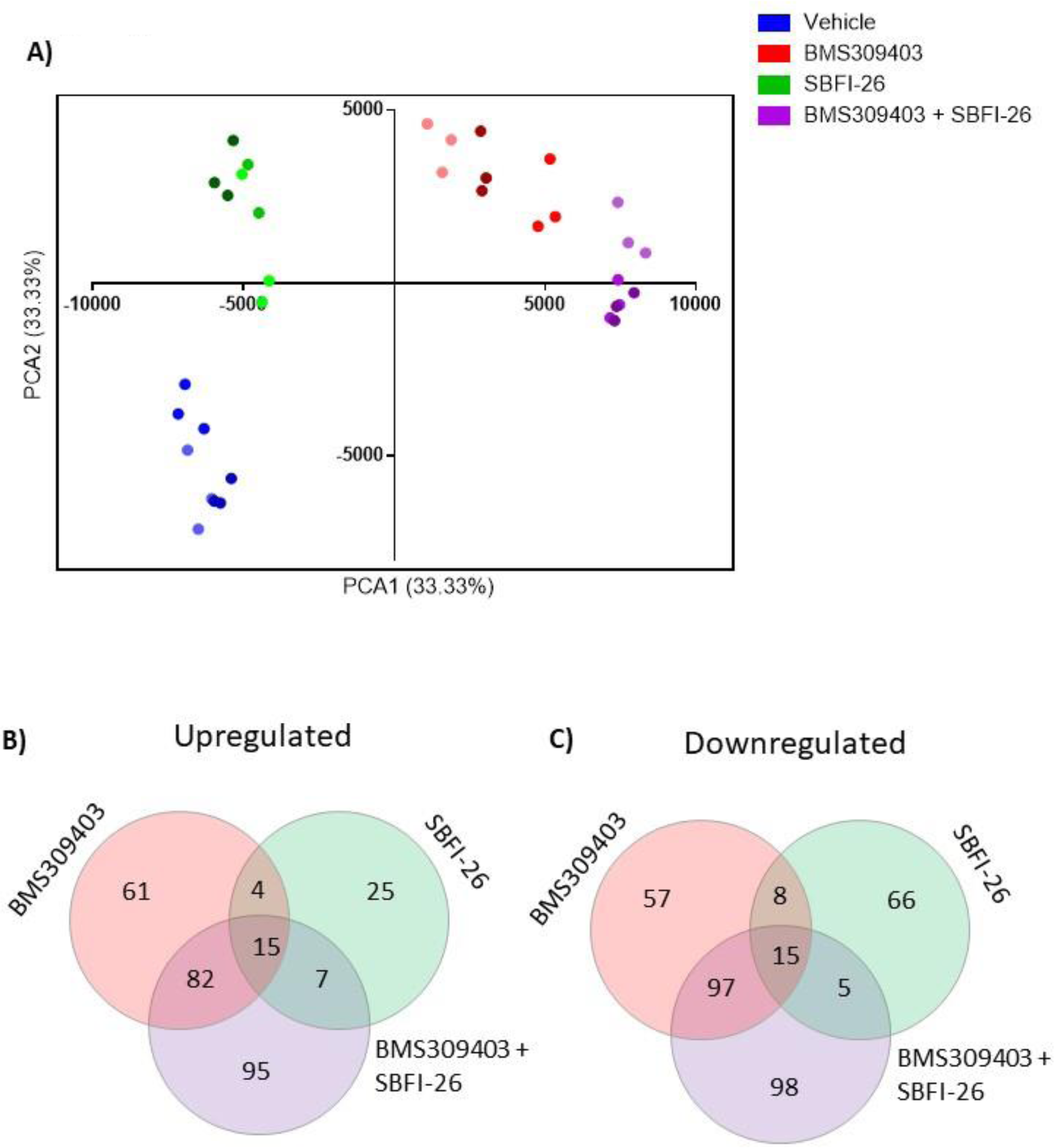
Mass spectrometry analysis revealed 48 hour treatment with FABP inhibitors induces a significant change in proteomic profile. A) Principal Component Analysis (PCA) of Proteomics data in MM.1S cells after 48 hour treatments with BMS309403 (50 µM), SBFI-26 (50 µM) or the combination, using Pareto Scaling. Shading of color indicates groupings of technical repeats within biological repeats. B) Venn diagrams of upregulated and C) downregulated proteins after treatment with FABPi. Data represents 3 biological replicates, with a significance cut off of p≤0.05 and a fold change of ±1.2.

**Supplemental Figure 12.**
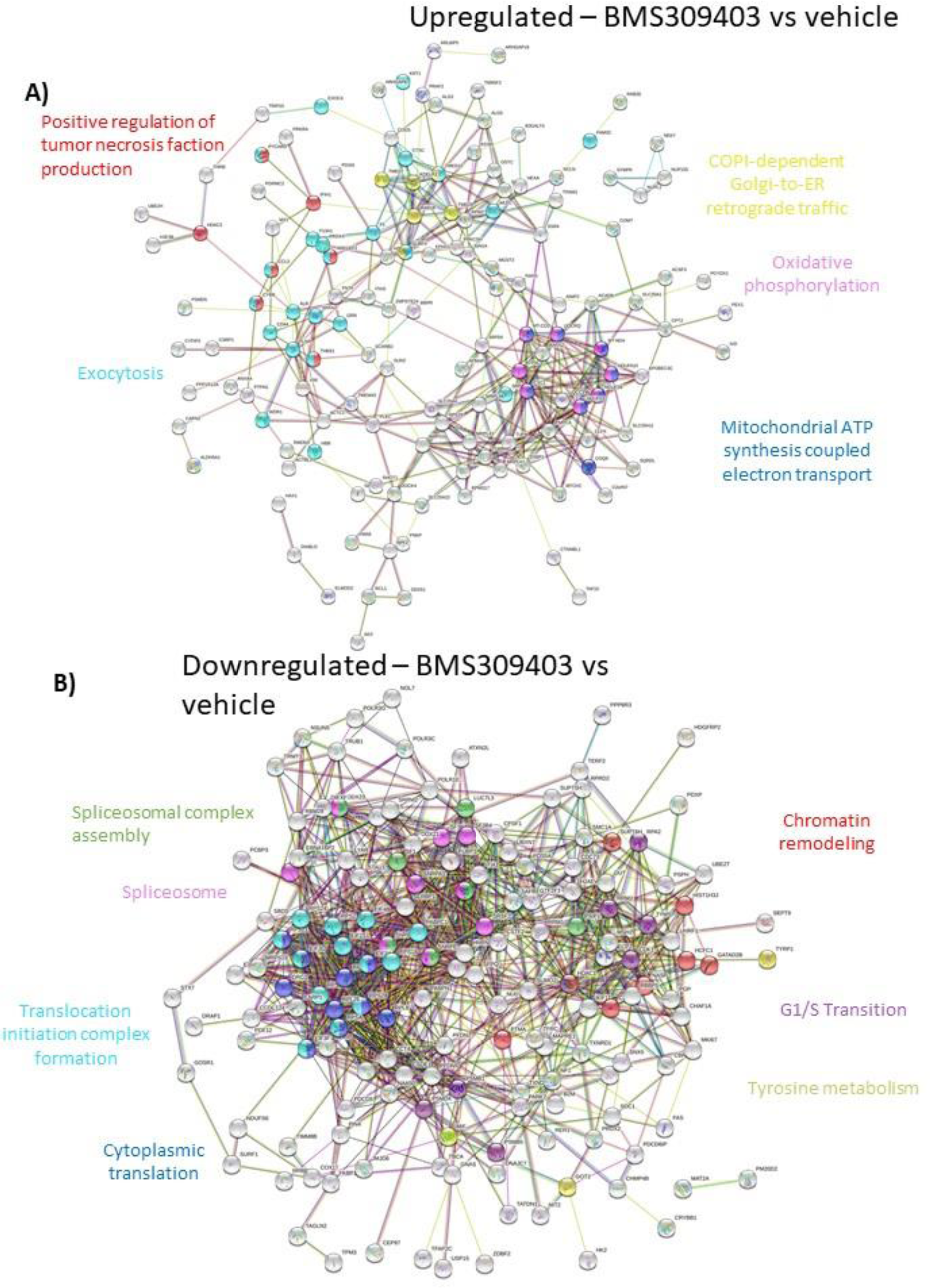
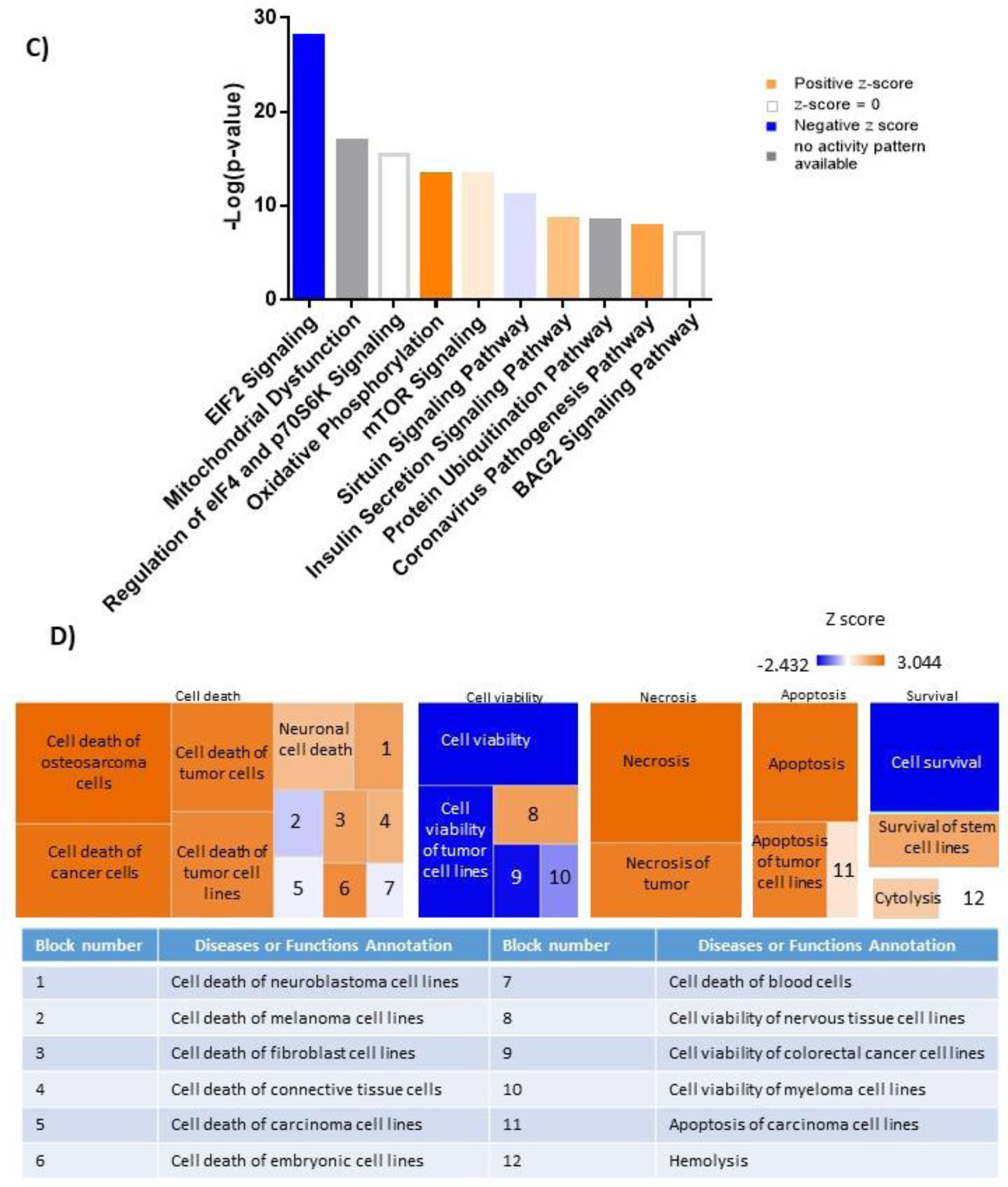
Mass spectrometry analysis reveals a shift in the proteomic profile MM.1S cells treated with BMS309403 for 48 hours. A) String db visualization of significantly upregulated and B) downregulated proteins with FABP inhibition. Color nodes represent the pathways implicated with treatment. C) Top 10 significantly changed pathways with FABP inhibition. For IPA analysis, orange represents positive z-score, blue indicates a negative z-score, gray represents no activity pattern detected and white represents a z-score of 0. D) Ingenuity pathway analysis revealed up- and downregulated pathways that correspond to the Cell Death and Survival heatmap. Z-score scale spans -2.432 to 3.044.

**Supplemental Figure 13.**
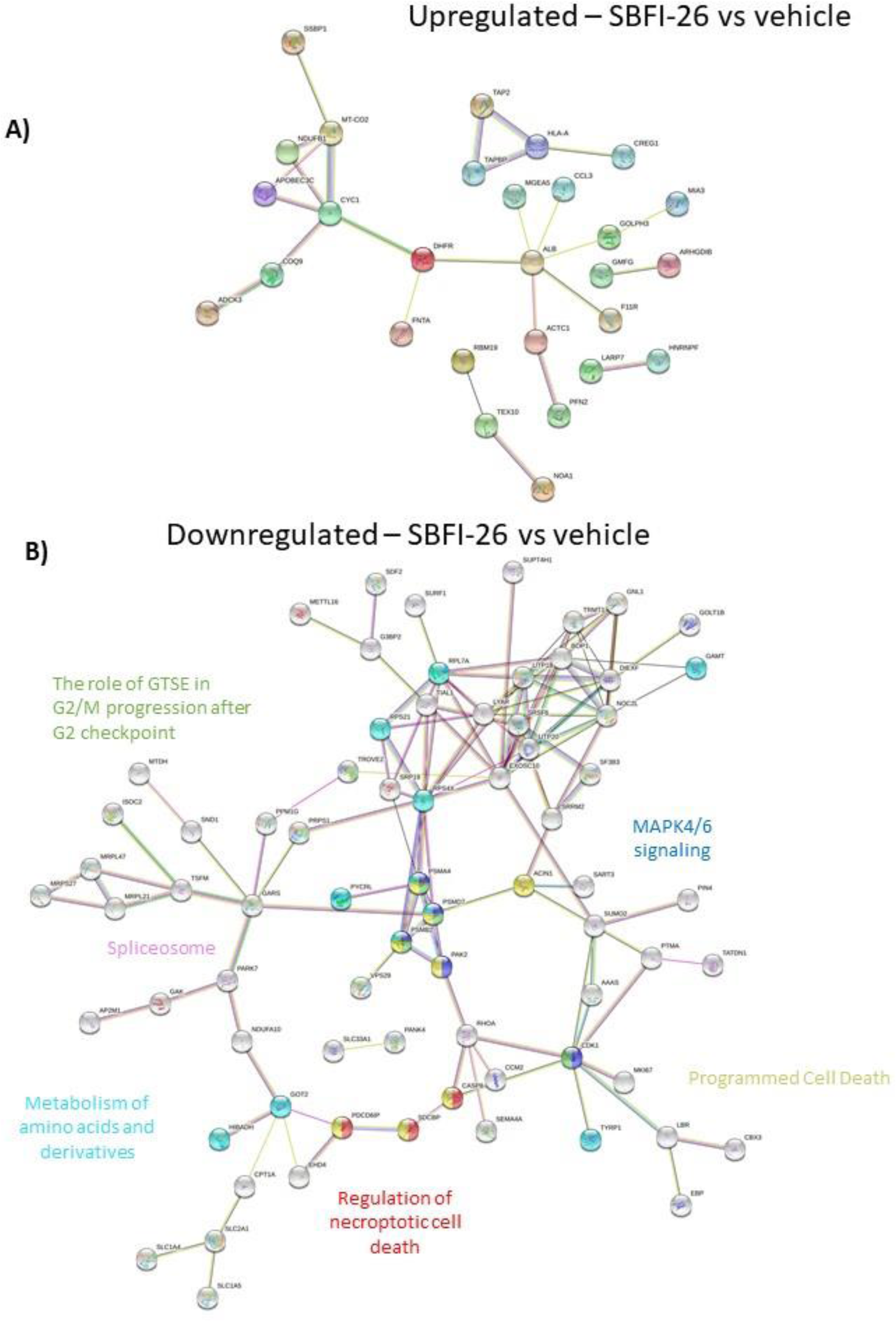
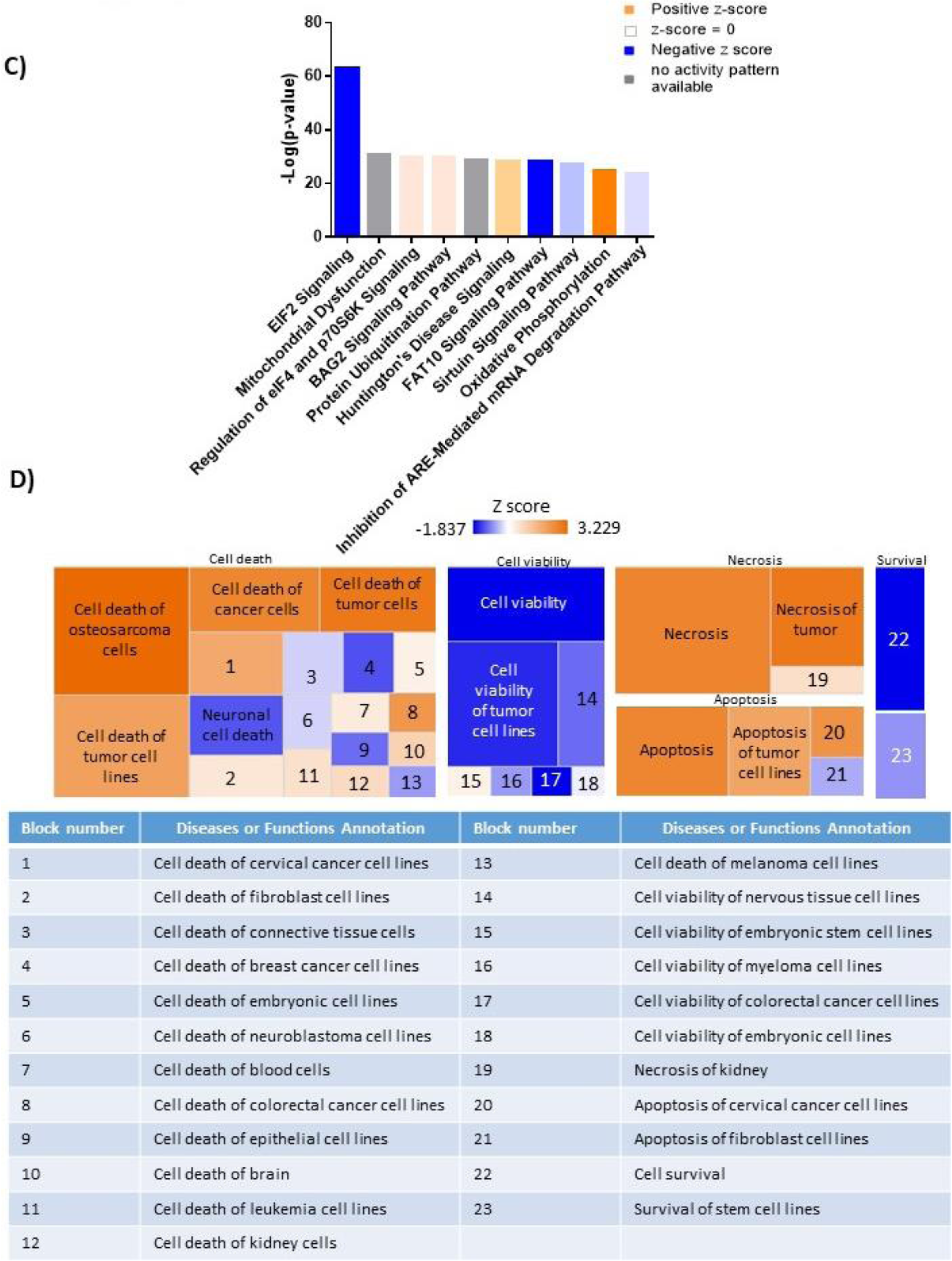
Mass spectrometry analysis reveals a shift in the proteomic profile MM.1S cells treated with SBFI-26 for 48 hours. A) String db visualization of significantly upregulated and B) downregulated proteins with FABP inhibition. Color nodes represent the pathways implicated with treatment. C) Top 10 significantly changed pathways with FABP inhibition. For IPA analysis, orange represents positive z-score, blue indicates a negative z-score, gray represents no activity pattern detected and white represents a z-score of 0. D) Ingenuity pathway analysis revealed up- and downregulated pathways that correspond to the Cell Death and Survival heatmap. Z-score scale spans -2.432 to 3.044.

**Supplemental Figure 14.**
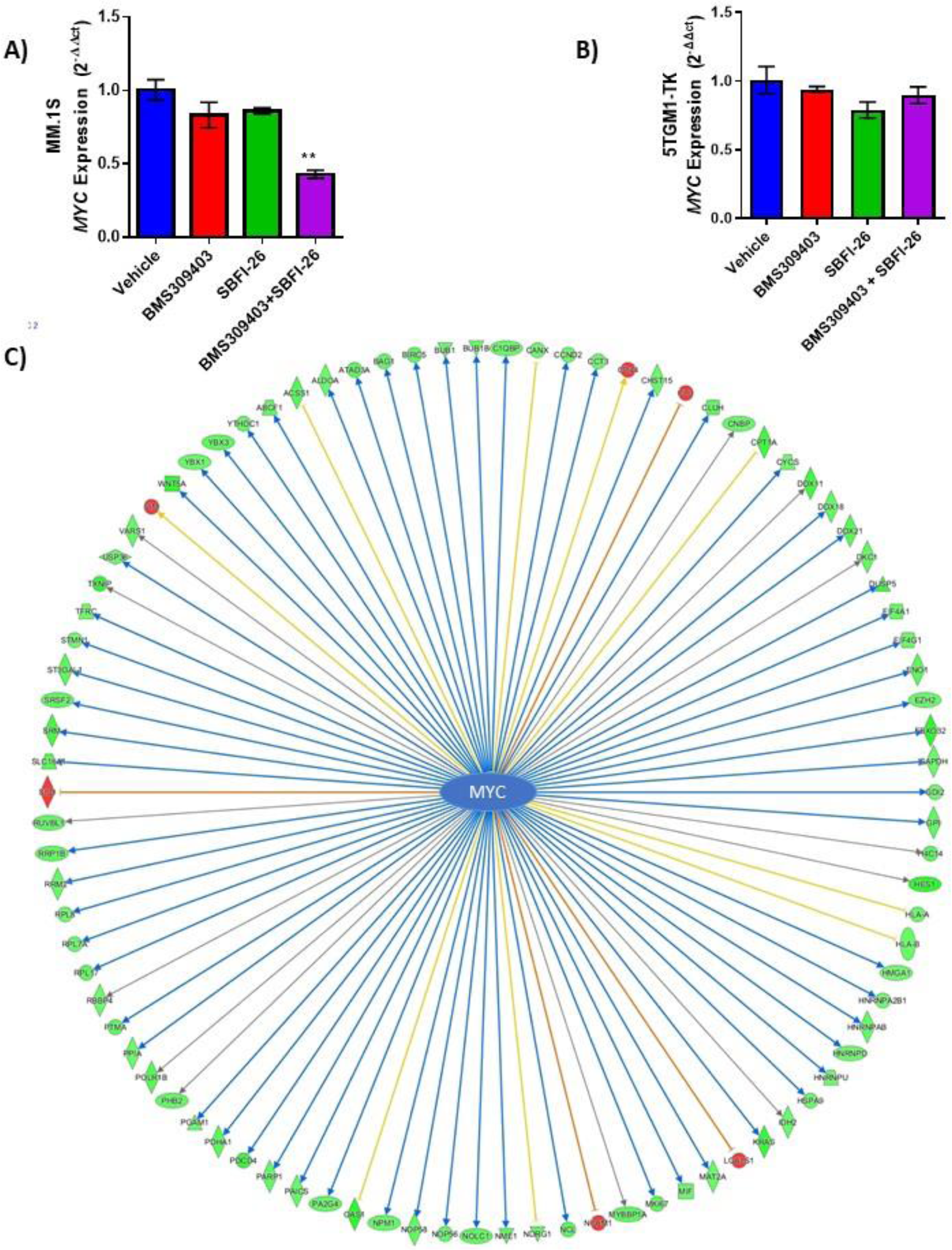
FABP inhibitor treatment alters expression of Myc gene expression and MYC-regulated genes. A) qRT-PCR of *MYC* in MM.1S and B) 5TGM1-TK cells after 24 hour treatments with BMS309403 (50 µM), SBFI-26 (50 µM) or the combination normalized to RPLP0 housekeeping gene. C) Co-treatment with BMS309403 and SBFI-26 induced changes in 91 genes modulated by MYC. Arrows indicate expression consistency with predicted patterns (blue=consistent, yellow=unknown, orange=inconsistent; color of molecules indicates expression pattern (green=decreased in co-treatment, red=increased in co-treatment). Data is plotted as mean ± SEM and analyzed with one-way ANOVA, **p<0.01, n=3.

**Supplemental Figure 15.**
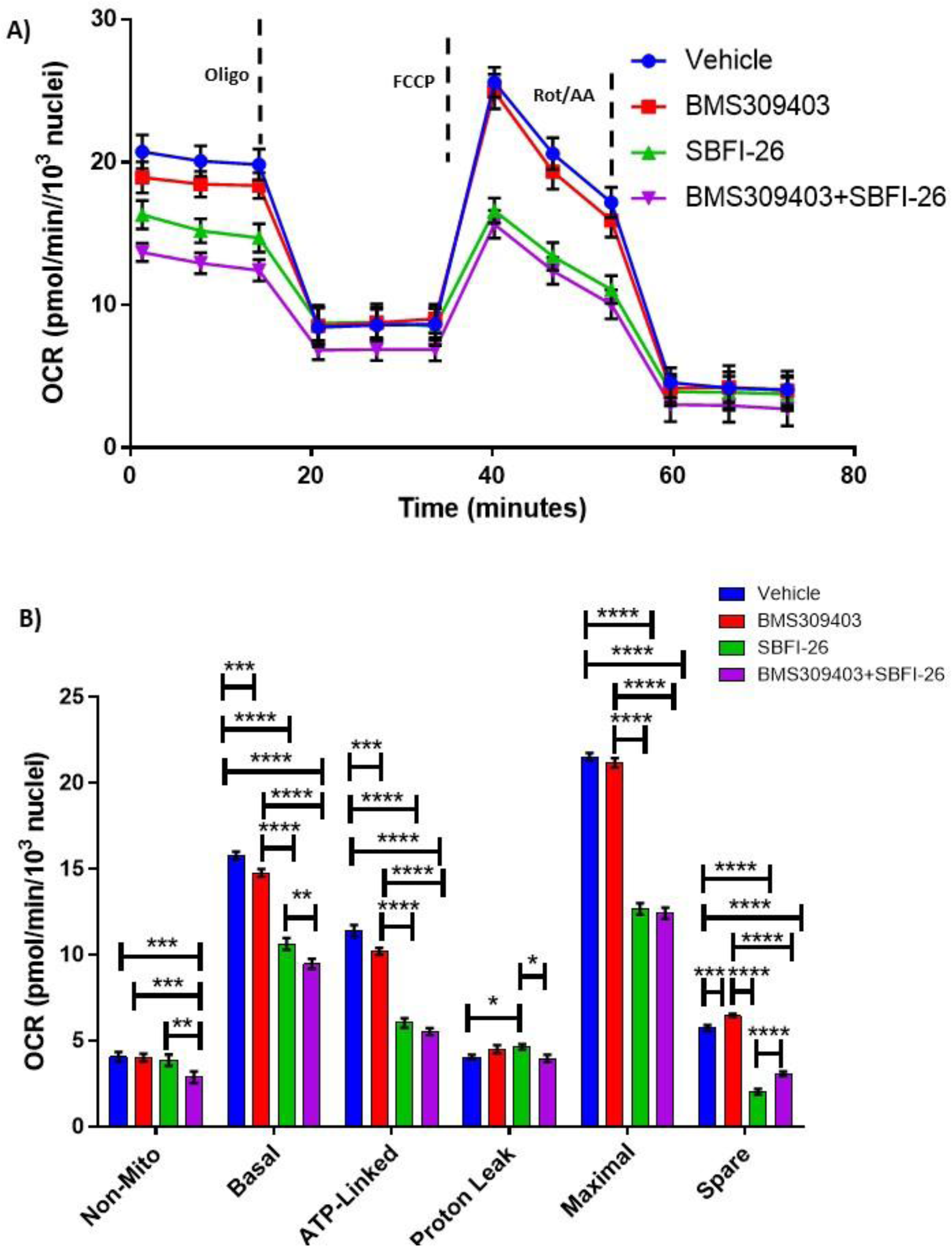
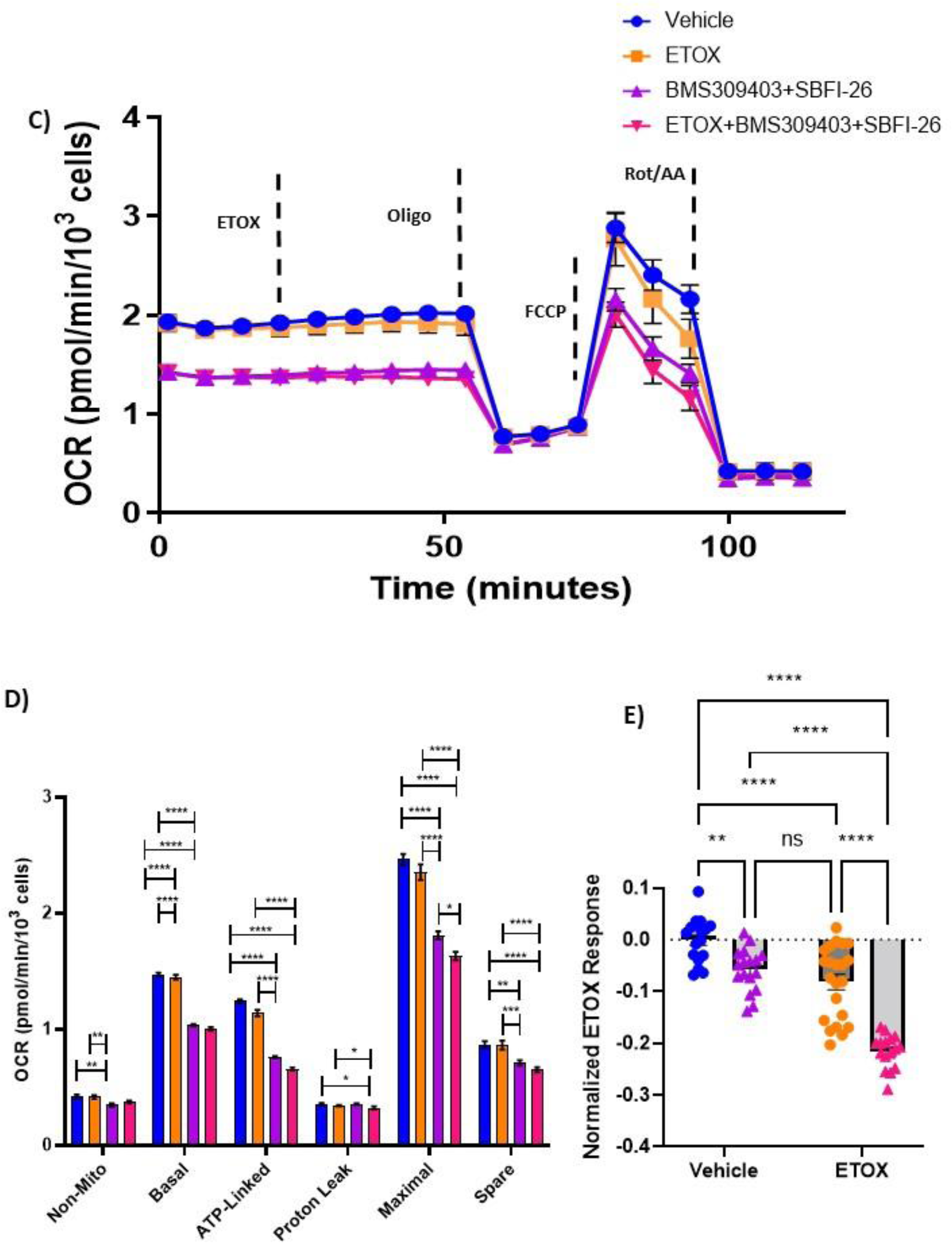
FABP blockade reduces cell metabolism and fatty acid oxidation: A) GFP^+^/Luc^+^MM.1S cells were cultured for 24 hours with BMS309403 (50 µM), SBFI-26 (50 µM), or both and then plated for Seahorse XF96 analysis in 96-well format. Oxygen consumption in cells was measured in basal conditions and in response to oligomycin (1.25 µM), FCCP (1 µM), and rotenone and antimycin A (.5 µM). Results represent 5 independent experiments with 1 representative experiment shown. In a separate set of experiments (n=2) cells were treated as above, however etomoxir or vehicle was added at a final concentration of 4µM prior to subjecting the cells to the mitochondrial stress test (C,D). ETOX response data normalized to MM.1S Vehicle control cells (E). Data represent Mean ± SEM; two-way ANOVA was used for each parameter with Uncorrected Fisher’s LSD multiple comparison post-hoc testing for significance shown as: *p < 0.05, **, p < 0.01, ***p < 0.005, ****p < 0.001. ns=non-significant.

**Supplemental Figure 16.**
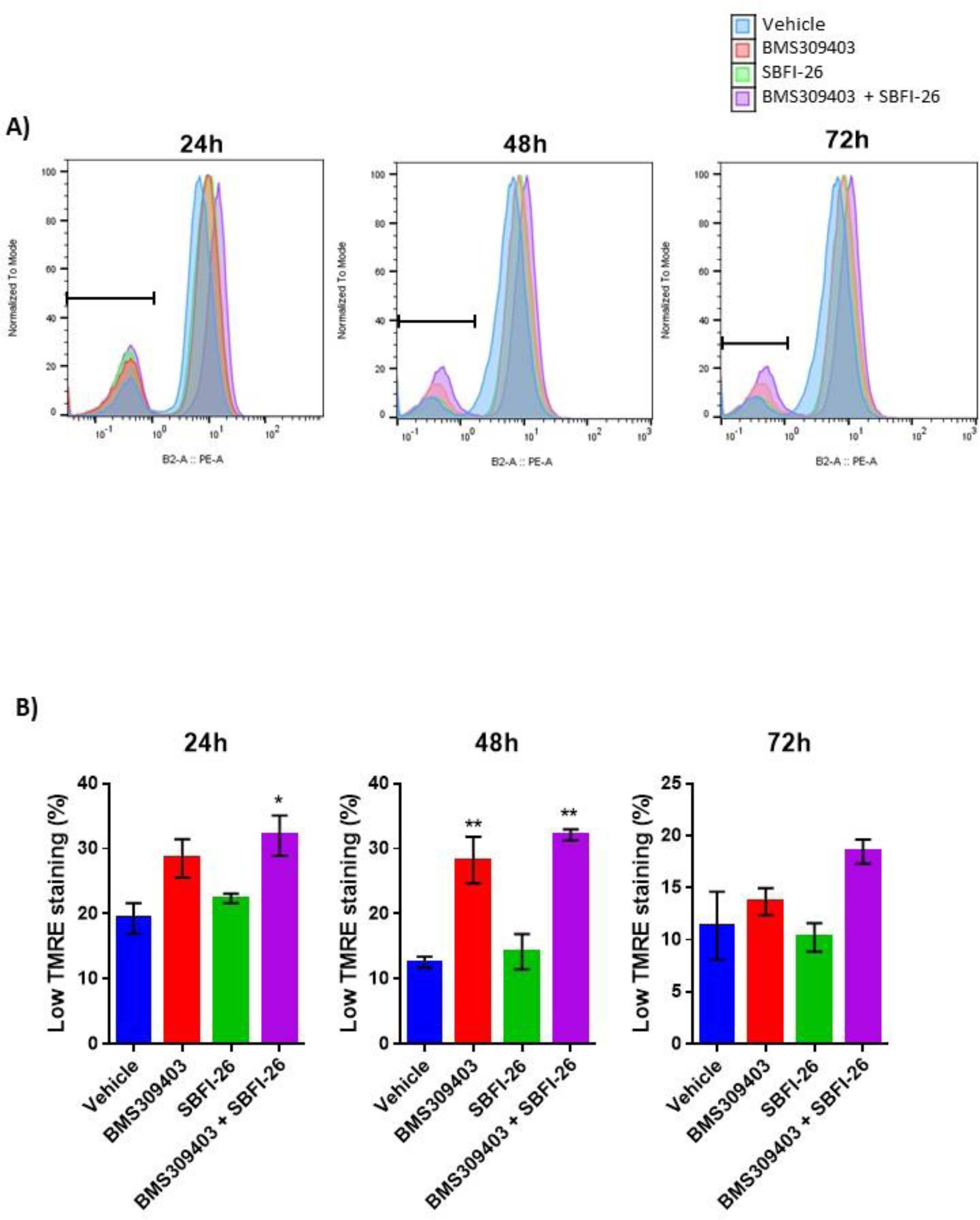
TMRE staining reveals compromised mitochondria in response to BMS309403 and the combination treatment. GFP^+^/Luc^+^MM.1S cells were treated with vehicle (DMSO), BMS309403, SBFI-26, or the combination treatment for 24, 48 or 72h prior to staining with tetramethylrhodamine, ethyl ester (TMRE). Representative TMRE staining and flow cytometry gating (A) demonstrate an increase in TMRE (low) stained cells with BMS309403 or the combination treatment relative to vehicle. Low TMRE stained cells are quantified in panel B; plotted as mean ± SEM and analyzed with one-way ANOVA, **p<0.01, n=3.

**Supplemental Figure 17.**
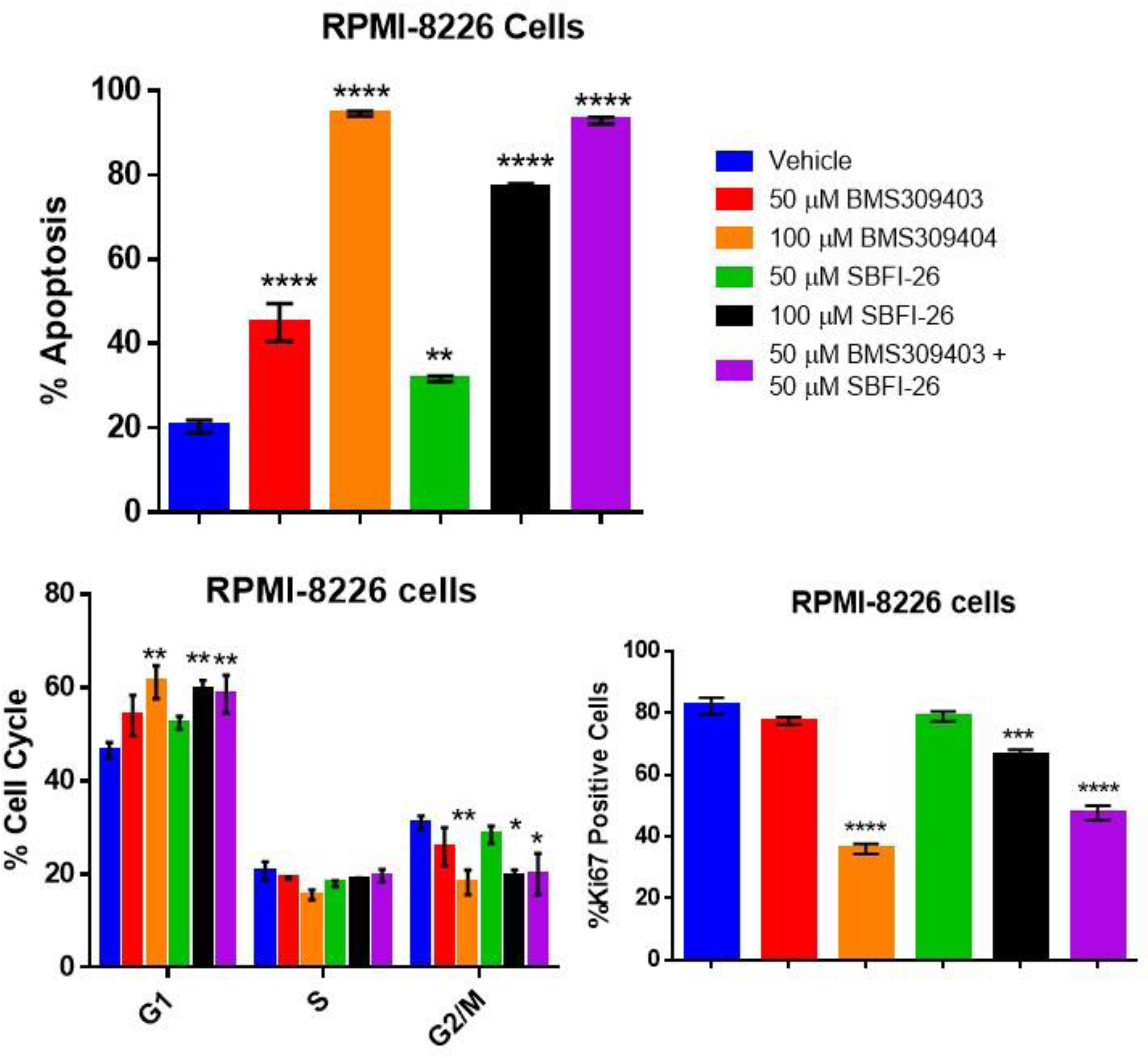
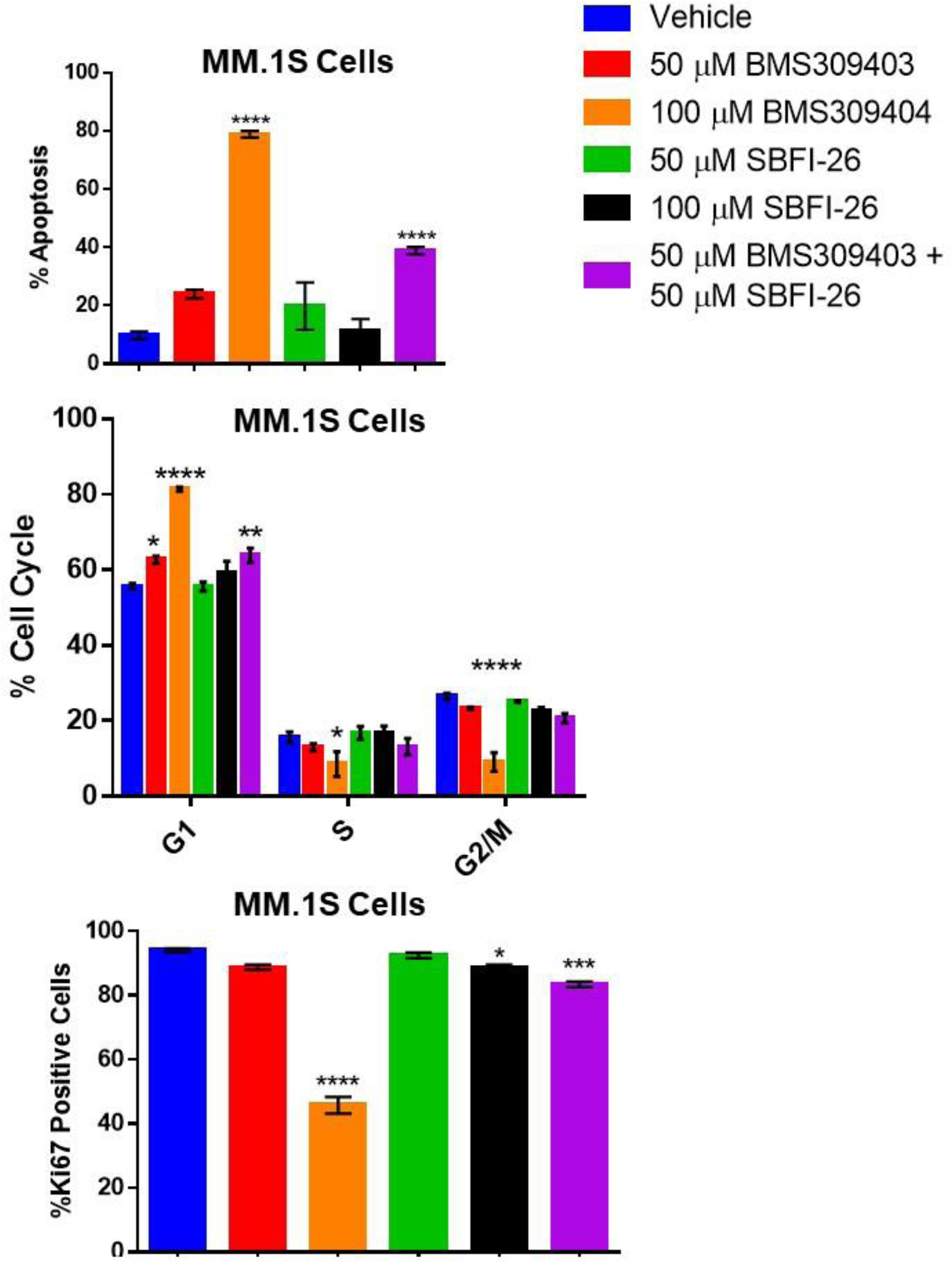
Apoptosis, cell cycle arrest and reduction in Ki67 positivity is induced in myeloma cells through inhibition of FABP proteins at 72 hours. Characterization of FABP inhibition on (A) RPMI-8226 cells and (B) GFP^+^/Luc^+^MM.1S cells.Data represent Mean ± SEM of least 3 experimental repeats with at least 2 technical samples per experiment. One-way ANOVA and Dunnett’s Multiple comparison testing was used; *p < 0.05, **, p < 0.01, ***p < 0.005, ****p < 0.001.

**Supplemental Figure 18.**
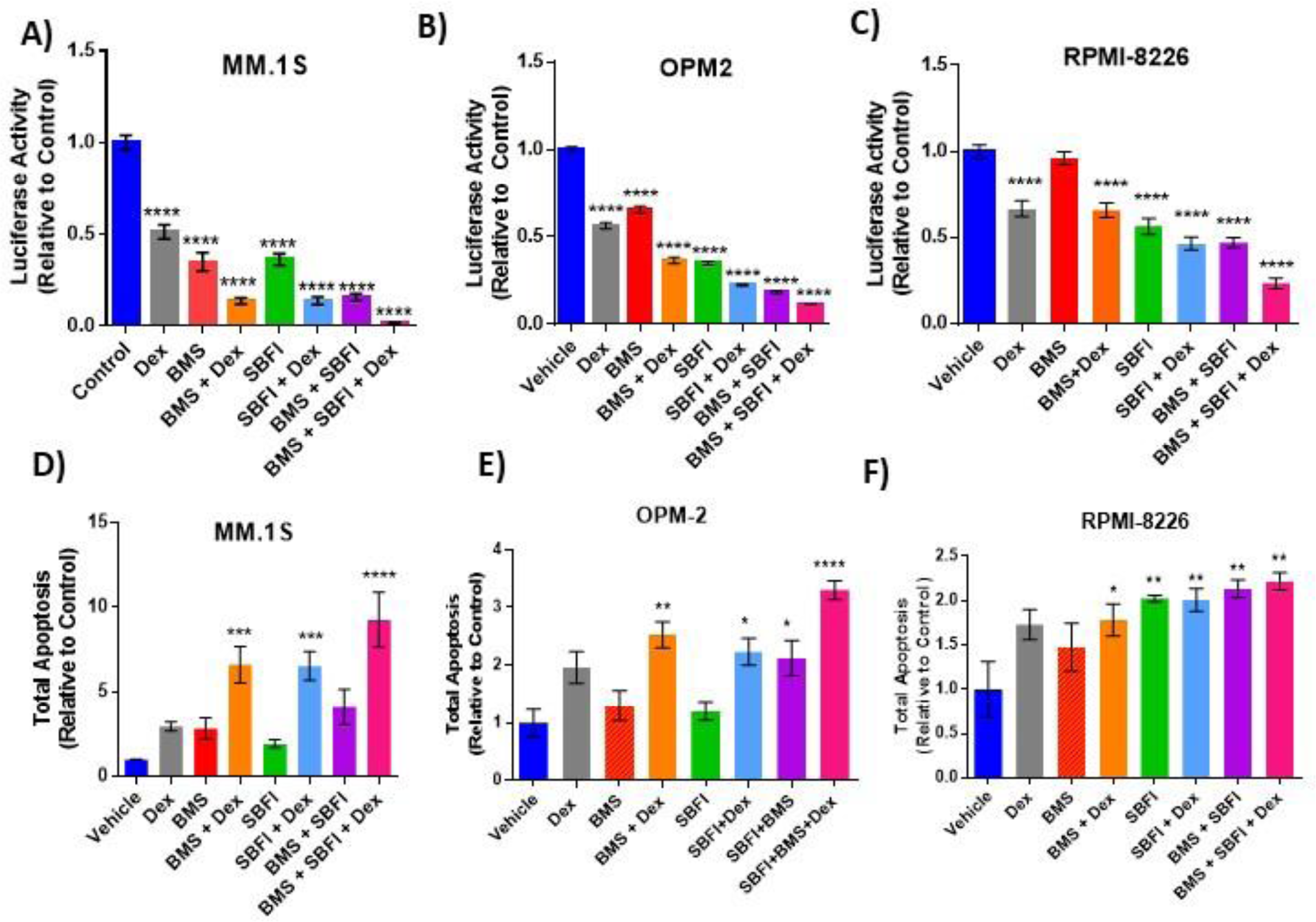
The effects of FABP inhibitors combined with dexamethasone after 72 hours *in vitro*. Combination of inhibitor treatment with dexamethasone results in significantly reduced cell numbers in MM.1S (A, n=3) and OPM-2 (B, n=4) as assessed by luciferin spike-in and RPMI-8226 (C, n=3) as assessed by CellTiter-Glo (normalized to control). Combination treatments also result in significantly elevated apoptosis in MM.1S (D, n=3), OPM-2 (E, n=3) and RPMI8226 (F, n=3). All graphs represent Mean ± SEM and significance compared to the control is determined by 1 way ANOVA with Dunnett’s multiple comparisons test. *p < 0.05, **, p < 0.01, ***p < 0.005, ****p < 0.001.

**Supplemental Figure 19.**
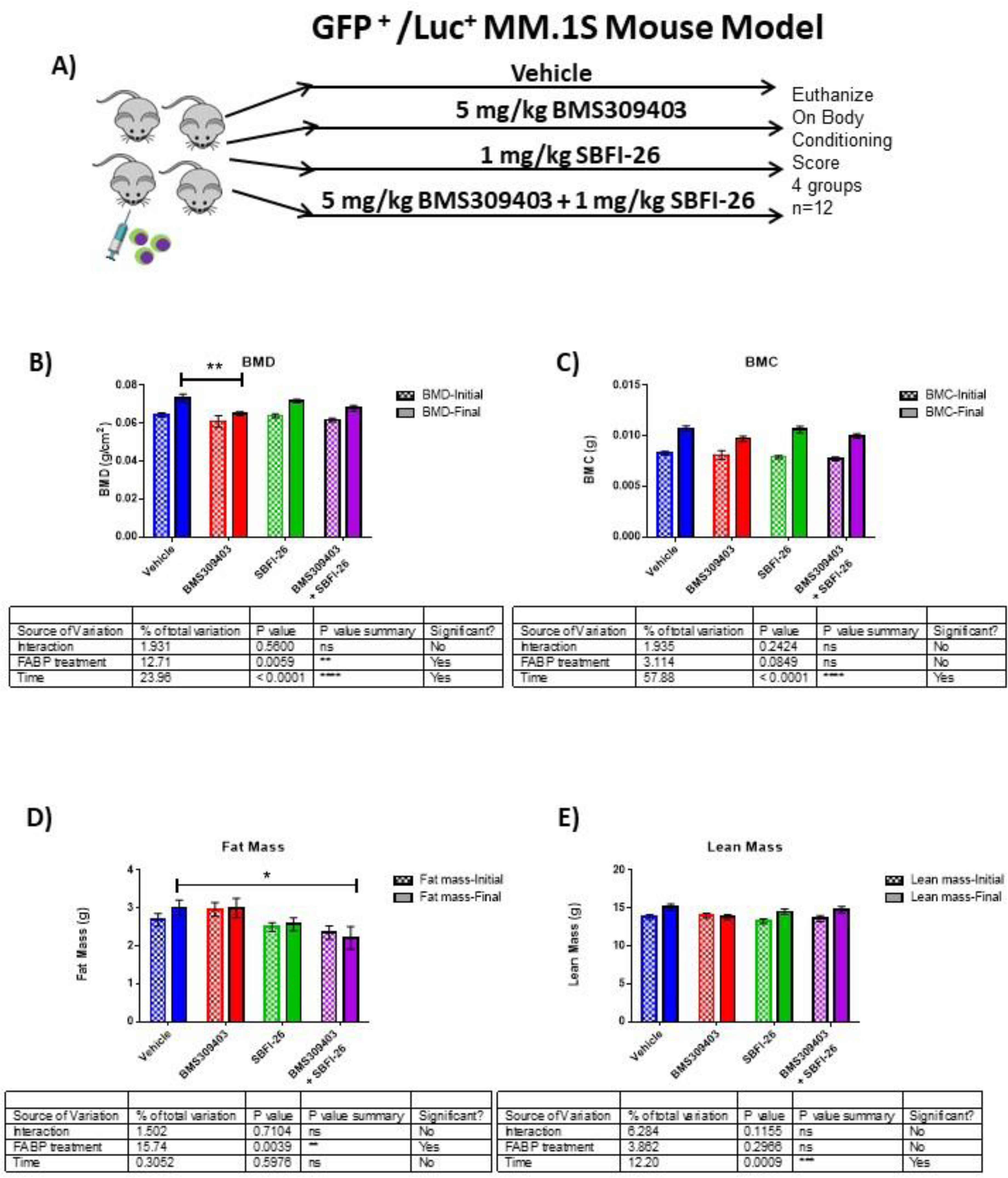
SCID-Beige-MM.1S^gfp+luc+^ *in vivo* characterization. A) *In vivo* model experimental flow chart. B-E) Piximus DEXA analysis at day 1 (Initial) and day 30 (final) after tumor cell inoculation plotted as mean ± SEM and analyzed with two-way ANOVA with Dunnett’s multiple comparison post-hoc test; *p<0.05, **p<0.01 n=8-12.

**Supplemental Figure 20.**
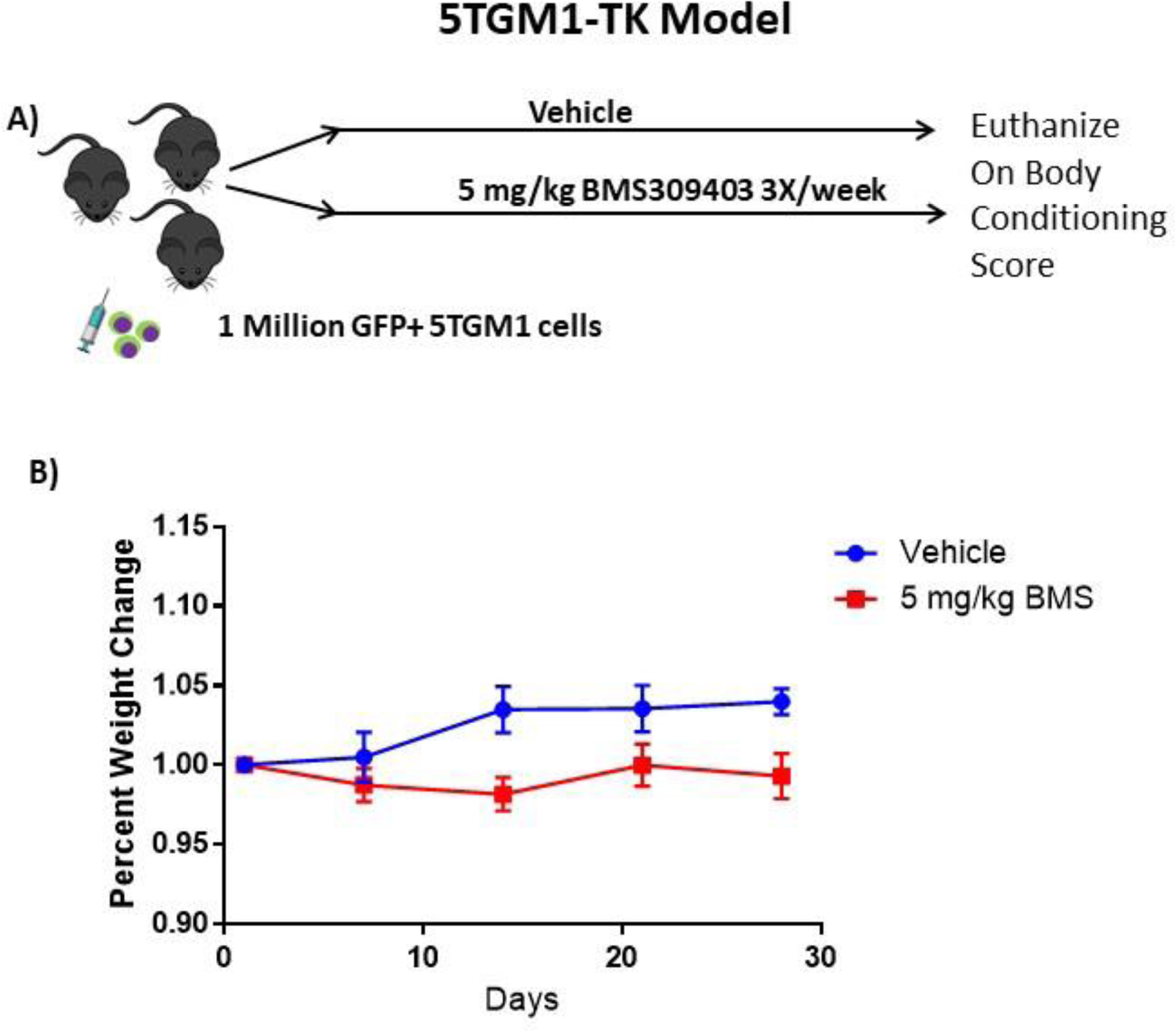
C57BL/KaLwRij-5TGM1^gfp+luc+^ *in vivo* characterization. A) *In vivo* model experimental flow chart. B) Mouse weights normalized to day 0 for each group, from day of injection plotted as mean ± SEM. Vehicle, n=8; BMS309403, n=9.

**Supplemental Figure 21.**
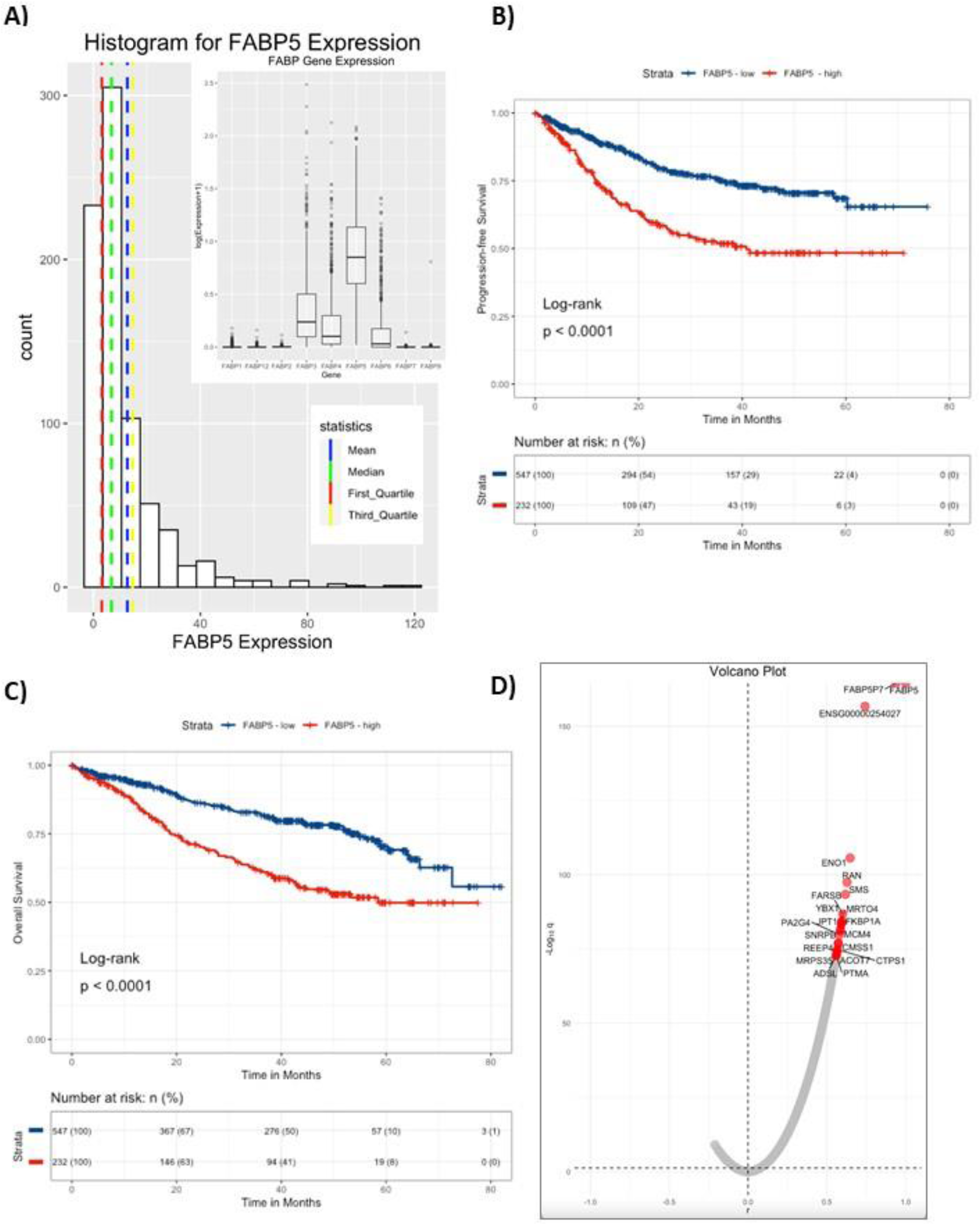
CoMMpass Dataset analysis of FABP5 demonstrates decreased progression-free survival and overall-survival in MM patients with high FABP5 expression in MM cells. A) Histogram of FABP5 in full dataset. Mean, Median, 1^st^ quartile and 3^rd^ quartile are marked. 779 cases total. Insert: Gene expression levels of all FABP genes in CoMMpass dataset. B) Progression-free survival was significantly shorter for MM patients with high expression of FABP5, with a hazard ratio of 1.64 (95% confidence interval 1.34–2.0). C) Overall survival was also significantly shorter for MM patients with high expression of FABP5, with a hazard ratio of 2.19 (95% confidence interval 1.66–2.88). D) Volcano plot of correlation analysis shows correlation between FABP5 and other genes in the CoMMpass dataset. A bias towards positively correlated genes was observed. Volcano plot was made by plotting the negative logarithm of the Benjamini-Hochberg adjusted p values (q) from a Pearson’s correlation test, which tested for correlation between FABP5 versus all other genes, with the top 20 genes highlighted in red.

**Supplemental Figure 22.**
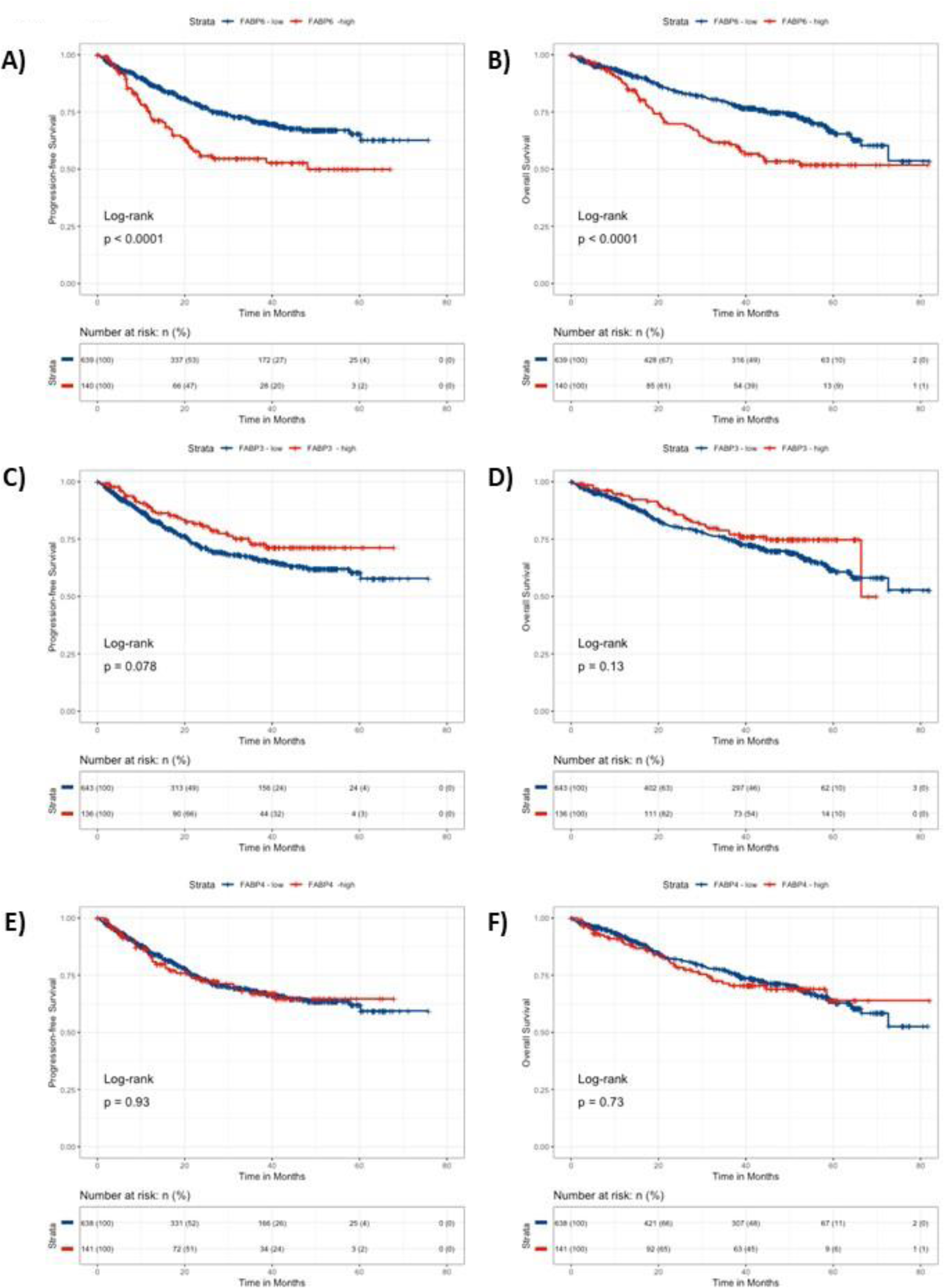
CoMMpass Dataset analysis of FABP4 and 6 demonstrates decreased survival and overall-survival in MM patients with high FABP6 expression in MM cells. A). 779 cases total. A) Progression free survival and B) Overall survival were significantly shorter for MM patients with high expression of FABP6, with a hazard ratios of 1.48 (95% confidence interval 1.172–1.869) and 1.837 (95% confidence interval 1.347–2.504) respectively. C) Progression-free survival and D) Overall survival for MM patients with high vs low expression of FABP3 were not significant. E) Progression-free survival and F) Overall survival for MM patients with high vs low expression of FABP4 were not significant.

